# Single cell analysis of FOXP3 deficiencies in humans and mice unmasks intrinsic and extrinsic CD4^+^ T cell perturbations

**DOI:** 10.1101/2020.07.06.189589

**Authors:** David Zemmour, Louis-Marie Charbonnier, Juliette Leon, Emmanuelle Six, Sevgi Keles, Marianne Delville, Safa Baris, Julien Zuber, Karin Chen, Benedicte Neven, Maria I Garcia-Lloret, Franck Ruemmele, Carlo Brugnara, Nadine Cerf-Bensussan, Frederic Rieux-Laucat, Marina Cavazzana, Isabelle André, Talal A. Chatila, Diane Mathis, Christophe Benoist

## Abstract

*FOXP3* deficiency in humans with IPEX syndrome and mice results in fatal systemic autoimmunity by altering regulatory T cell (Treg) physiology, but actual cellular and molecular mechanisms of disease are unclear, part because Treg surface markers may be unreliable in disease states. We used deep profiling by flow cytometry, population and single-cell RNAseq to analyze Tregs and conventional (Tconv) CD4+ T lymphocytes in cohorts of IPEX patients with a range of genetic lesions, and in *Foxp3*-deficient mice. In all patients and mice, heterogeneous Treg-like cells with an active *FOXP3* locus were observed, some differing very little from normal Tregs, others more distant. Tconv showed no widespread activation or Th bias. The dominant mark was a monomorphic signature equally affecting all CD4^+^ T cells, unexpectedly dampening tumor-Treg and cytokine-signaling modules. In mixed bone marrow chimeras, WT Tregs exerted dominant suppression, normalizing the states of mutant Treg and Tconv, extinguishing the disease signature, and revealing a small gene cluster truly regulated, cell-intrinsically, by FOXP3. These results suggest a two-step pathogenesis model, with therapeutic implications: limited downregulation of a few core Treg genes de-represses a systemic mediator(s), which imprints the disease signature on all T cells, and further dampens Treg function.

## INTRODUCTION

T regulatory cells (Treg) that express the transcription factor FOXP3 are dominant negative regulators of many facets of the immune system, controlling immune responses and enforcing peripheral tolerance to self, symbiotic commensals and fetal antigens ^1, 2^. They also have extra-immunological roles in maintaining tissue homeostasis outside of the purely immunological context ^3^. Tregs share a common transcriptional signature, genes differentially expressed relative to their conventional CD4^+^ counterparts (Tconv), in mice and humans ^4–8^. Much of the Treg signature is controlled by FOXP3, the lineage’s defining transcription factor (TF), although FOXP3 is neither fully necessary nor sufficient to establish the Treg transcriptome, requiring input from transcriptional cofactors ^9–14^. Consistent with their pleiotropic functions, Tregs show a range of phenotypic variation, differing by activation state, effector capabilities, and tissue localization ^3, 15^. Treg heterogeneity has been further refined by single-cell transcriptomics (scRNAseq) ^7, 16–18^.

IPEX (Immune dysregulation-Polyendocrinopathy-Enteropathy-X-linked) is perhaps the prototype of monogenic autoimmune syndromes ^19^, resulting from FOXP3 loss of function (LOF) ^20–23^ and thus Treg deficiency ^24–27^. It usually presents in early infancy and includes a constellation of autoimmune manifestations dominated by severe enteropathy, eczematous dermatitis and type-1 diabetes, with less common manifestations (nephropathy, hypoparathyroidism, antibody-mediated cytopenias) ^28–31^. There are several causes to the variable severity observed in IPEX patients. First, complete LOF mutations are generally more deleterious than missense mutations; and mutations in the DNA-binding forkhead or dimerization (leucine zipper) domains are generally more severe than N-terminal mutations, in keeping with results from our alanine-scan dissection of FOXP3 ^14, 30, 32, 33^. But manifestations and severity can vary between patients with the same mutation ^30, 34^, suggesting that genetic modifiers, environmental influences or immunological history modify the disease course in each patient. Mice of the spontaneous *scurfy* mutant line, or carrying engineered *Foxp3* LOFs, show a similar rapidly lethal phenotype, dominated by enteropathy, dermatitis and lymphoproliferation ^1, 35^; partial or slower disease appears in mice with mild *Foxp3* missense mutations ^14, 36–38^.

Also contributing to this variability, the relationship between FOXP3 and Treg cells is now recognized not to be obligate. Several reports have documented the existence of Treg-like cells in the absence of viable *Foxp3* in *scurfy* mice ^39–41^ and some IPEX patients ^42–44^. In addition, FOXP3 is not exclusive to Treg cells: it is also expressed, albeit at lower levels than in Tregs, early after Tconv activation ^45–48^, and scRNAseq showed some FOXP3-positive cells otherwise similar to Tconv cells ^7^.

The advent of single-cell transcriptomics opened the potential to revisit the alterations of CD4^+^ T cells in IPEX patients, which were previously difficult to address since the markers potentially used to sort Treg or Tconv cells may be themselves perturbed. We have thus performed a deep profiling analysis in IPEX patients that associates flow cytometry, single-cell (for resolution) and conventional (for depth) RNAseq. Complementary analyses in *scurfy* mice assessed the generality of the conclusions, eschewing the inevitable confounders of patient material, and permitting experimental manipulations to trace the source of the perturbations. The results paint an unexpected landscape of IPEX T cells, with a complex mix of Treg-like cells, some healthy-like and others more perturbed, a surprisingly narrow intrinsic signature of FOXP3 but an unexpectedly dominant “IPEX signature” that affects both Treg and Tconv cells, and suggests a feed-forward loop in T cell perturbations culminating in clinical disease.

## RESULTS

### Study Cohorts

This study included a primary and a replication cohort for confirmation and refinement (including some re-sampling of initial patients). Altogether, we analyzed peripheral blood mononuclear cells (PBMC) from 15 IPEX patients and 15 healthy donors (HD) collected at two reference clinical centers (Necker Hospital, Paris and Children’s Hospital, Boston; Table S1, Fig. 1a). HD had no significant personal medical history and were recruited during well-child visits or orthopedic follow-ups. IPEX presentation was typical, first symptoms appearing neonatally for most (1-8 weeks). As in other cohorts ^28–31^, enteropathy (12/14), dermatitis (10/14), food allergy (8/13) and diabetes (6/14) were most common, with less frequent kidney, neurological or pulmonary involvement. Four patients were untreated at the time of sampling, but most were managed by immunosuppression (mainly mTOR inhibitors) – four of them later received bone marrow transplants. Blood samples were collected at different ages (9 months to 26 years) during routine visits, with no concurrent acute events. Five patients were analyzed at two or three time points (6-12 months apart) to assess the stability of the transcriptional characteristics.

**Fig. 1.**
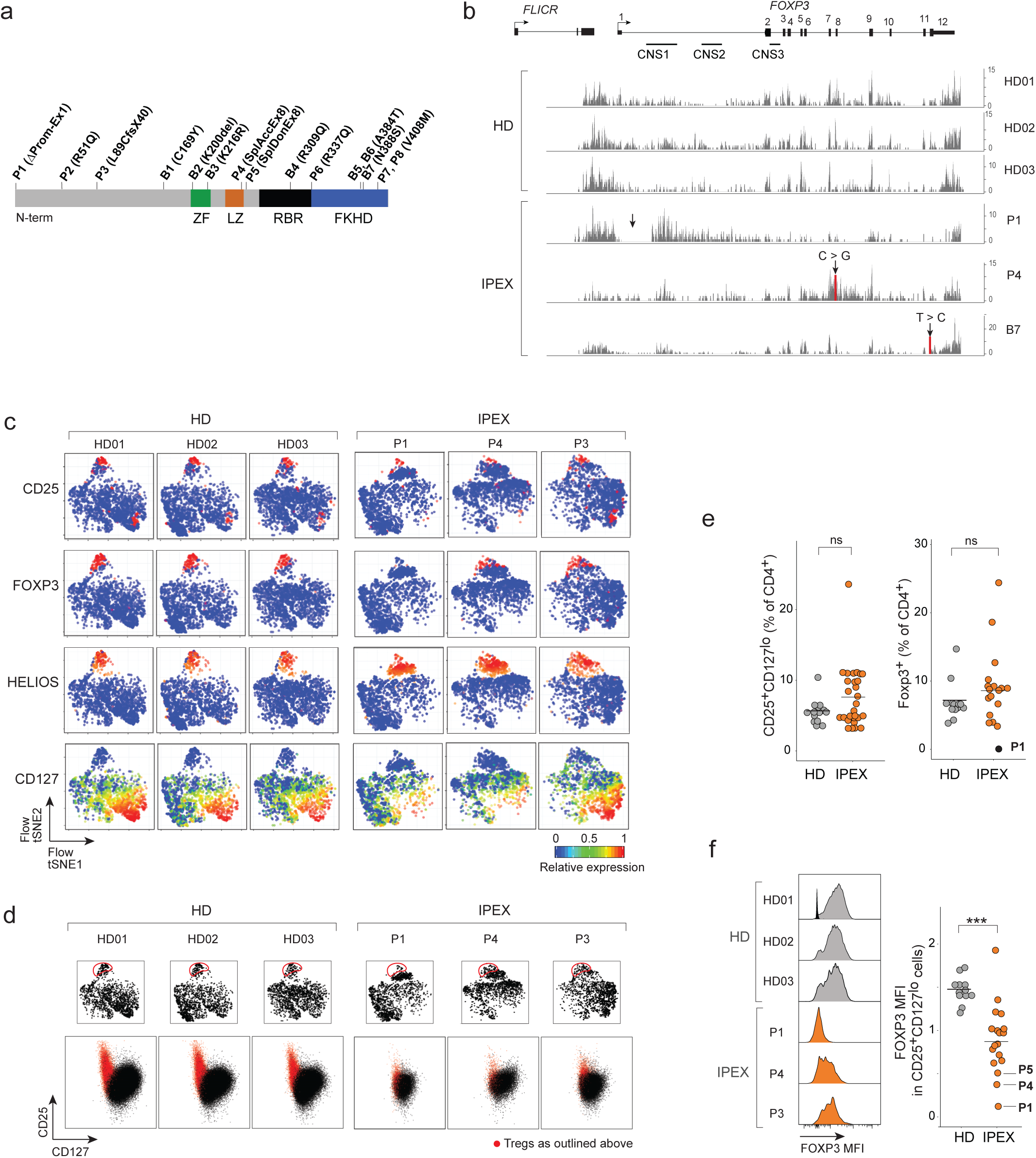
Identification of Treg-like cells in IPEX by flow cytometry. **a.** Position of the FOXP3 mutations in the IPEX cohorts (domains: ZF: zinc-finger; LZ: leucine-zipper; RBR: RUNX1 binding region; FKH: Forkhead). **b.** Mapping of RNAseq reads from CD4^+^CD25^+^CD127^lo^ cells to the *FOXP3* locus in representative samples (see Fig. S1). Arrow indicates mutation. **c.** Flow-tSNE plot of CD4+ T cells, from anti-CD3, CD4, CD25, CD127, HELIOS, CD45RA and FOXP3 staining. Color represents scaled expression. **d.** Flow cytometric anti-CD25/CD127 plots of CD4+ T cells (bottom row); red: cells gated in the flow-tSNE plot, top row. **e.** Proportion of CD25^+^CD127^lo^ cells and total FOXP3^+^ cells in HD and IPEX (each dot a sample). **f.** FOXP3 expression in CD25^+^CD127^lo^ HD and IPEX cells. Left: representative cytometry profiles (see Fig. S1 for all other samples; one unstained control overlaid with HD01), quantitation at right (each dot a sample; *** t.test p < 0.0001).

*FOXP3* mutations were confirmed by sequencing. Most were missense mutations in different domains, two of them represented twice (V408M and the common A384T). One mutation (P1) was a large deletion spanning the promoter and intron 1; two others affected the exon 8 splice acceptor and donor sites (P4 and P5, respectively). The mutations’ impact was seen in the RNAseq reads at the *FOXP3* locus of sorted CD4^+^CD25^hi^CD127^lo^ cells (see below). For HD, the traces were very reproducible, reads piling up at the exons, the conserved non-coding sequence (CNS) 1 and the lncRNA *FLICR* (Fig. 1b, S1). Profiles from patients were variable: generally normal for the missense mutations; for P1, aberrantly initiated transcripts piled up around CNS1 with essentially no exonic reads; for P4, the splice mutation in exon 8 disturbed later exons and introns. Importantly, though, the *FOXP3* locus was active in all IPEX cells, at far greater levels than in Tconv (Fig. S1). Interestingly, FLICR transcripts varied markedly in patient cells, unrelated to the proportion of *FOXP3* exonic reads, in line with its independent transcriptional regulation ^49^.

### Identification of Treg-like cells in IPEX patients

Tregs are typically identified by flow cytometry as CD25^hi^CD127^lo^, a combination that overlaps well with intracellular FOXP3 expression ^50^. Such cells have been reported in IPEX patients ^42–44^, but we first adopted a more global approach to identify Treg-like cells, since expression of these two markers could be perturbed by FOXP3 deficiencies. We analyzed CD4^+^ PBMCs by multiparameter flow cytometry for a larger panel of Treg markers (CD25, CD127, CD45RA, CD3 and CD4 - whose expression is subtly lower in Treg vs Tconv), coupled with intracellular staining for FOXP3 and HELIOS (Fig. S2). The data from individual staining experiments, including HD and IPEX samples, were then integrated in 2D space by t-Distributed Stochastic Neighbor Embedding (t-SNE; Fig. 1c, Fig. S3). Tregs from HD donors clustered tightly, with characteristic FOXP3, CD25 and HELIOS expression. For IPEX donors, the same region of the plots contained cells that also expressed CD25 and HELIOS, but variable FOXP3 (predictably absent for P1, more normal for P4). Indeed, “gating” Treg-like cells on the tSNE showed that they fell in the expected position of the CD25/CD127 plot (Figs. 1d, S2a), although with lower CD25 in IPEX samples. The proportion of Treg-like cells, and generally of FOXP3+ cells, were well conserved in IPEX patients (Fig. 1e, Fig. S2b), although the levels of FOXP3 protein were variably reduced (Fig. 1f). Other than the low or null levels resulting from promoter or splicing mutations, there was no discernible relationship between FOXP3 intensity and the domains affected by each mutation. These cytometry results confirmed that cells with surface Treg characteristics, but with low or absent FOXP3, could be identified in every one of the IPEX patients.

### Widespread perturbations of IPEX CD4+ T cells transcriptomes

Given the presence of Treg-like CD25^hi^CD127^lo^ cells in PBMCs from IPEX patients, we performed population low-input RNAseq after sorting these and Tconv (CD25^-^CD127^hi^) cells, as a preliminary to single-cell profiling and to help anchor its interpretation (first and replication cohorts profiled independently). Several observations emerged. First, the classic Treg signature was well conserved in Tregs from IPEX donors overall (Fig. 2a), including prototype transcripts (*IL2RA, LRRC32, CTLA4*). In ranked FoldChange (FC) plots that display the Treg signature for individual donors, the distribution of signature genes was tight for HD but more variable in IPEX Tregs, ranging from quasi-normal (P7) to markedly affected (P6, P8; Fig. 2b, S4a) – perhaps surprisingly, P1 with the complete LOF mutation did not show the strongest reduction in Treg signature genes. Interestingly, the Treg signature intensity score seemed better preserved in Tregs from IPEX donors than its coefficient of variation (Fig. 2b,c), suggesting an instability of the Treg signature in the absence of a fully functional FOXP3. These indices did not correlate with clinical outcomes (Fig. S4b). The IPEX mutations did not affect all signature genes equally, however: some Treg signature genes were actually over-expressed in IPEX Tregs (*DUSP4, LRRC32, CTLA4*), while most were downregulated as expected (Fig. S4c).

**Fig. 2.**
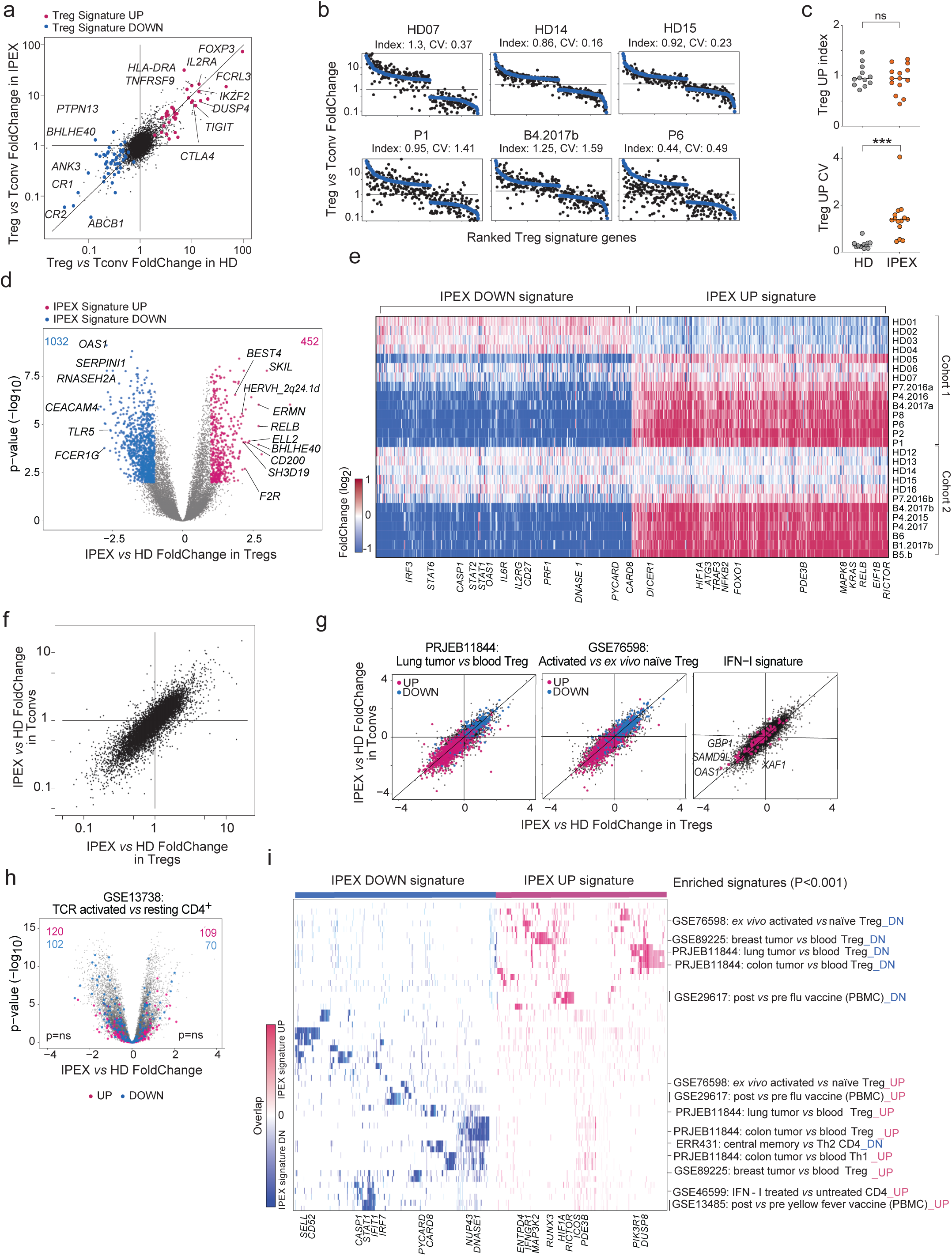
Transcriptional changes in IPEX Treg and Tconv cells by population RNAseq. **a.** Population RNAseq was performed on sorted Tconv and Treg-like cells from HD and IPEX donors. The Treg/Tconv FoldChange in HD (x-axis) and IPEX (y-axis); Treg signature ^6^ genes are highlighted. **b.** Ranked FC plots of Treg signature transcripts for individual donors, ranked according to mean FC in all HD (blue dots). FC values for each donor (black dots) computed from the donor’s Treg vs the mean of HD Tconv. **c.** Index and coefficient of variation (CV) of Treg Up signature transcripts (each dot is a sample). *** t.test p < 0.001. **d.** FoldChange vs p value (volcano) plot comparing normalized expression in all IPEX to all HD samples. Genes with differential expression (at p<0.01, FC >2) are highlighted, and numbers shown. **e.** Heatmap of the expression ratio, for IPEX signature genes defined in d, in Tregs from each donor, computed against the mean expression in HD Tregs (each cohort computed against its own HD Treg set). **f.** Comparison of mean IPEX effect (all IPEX vs all HD samples) in Treg (x-axis) versus Tconv (y-axis). **g.** Same FC/FC plot as in f, but highlighted with representative signatures of tumor-infitrating Tregs, activated Treg vs resting Treg, and IFN-I induced genes. **h.** Same volcano plot as in d, but with highlights from a representative CD4+ T activation signature (significance from hypergeometric test). **i.** Heatmap, for the IPEX signature genes defined in d, of their overlap with the pathways and signatures that significantly overlap with the IPEX signature (hypergeometric test p<0.001). Present genes are shown by tick marks, color-coded by their IPEX/HD FoldChange.

It is worth mentioning that no patient’s Treg or Tconv cells showed unusual expression of *IL4* or *IL5*, transcripts which we and others found to be paradoxically upregulated in response to forced expression of mutant FOXP3 ^14, 37^. More generally, there was no specific induction of cytokine genes in Tconv from IPEX patients that might have denoted a loss of Treg control (Fig. S4d).

We then assessed more generally the impact of IPEX mutations on transcriptomes of Treg-like cells. Widespread differences were observed (Fig. 2d), reproducibly in the two cohorts (Fig. S5a, Table S2). This “IPEX signature” was consistent in every patient, involving all the same genes, albeit at variable overall intensity (Fig. 2e). It did not correlate with the Treg signature intensity score (Fig. S5b) and was strikingly similar in Treg and Tconv cells (Fig. 2f), indicating a global impact on CD4^+^ T cells that transcended the sole effect of FOXP3 in Tregs (only 17 genes of the Treg signature belonged to this IPEX signature, Table S2). There was no marked relationship between the main clinical parameters, including current corticosteroid or rapamycin treatment, and the IPEX signature score (Fig. S5c), which was also present in untreated IPEX patients. The IPEX signature was not accompanied by activation of endogenous retroviruses, which might plausibly be reactivated (Fig. S5d). Enrichment analysis showed significant overlap between the IPEX signature and several gene expression signatures of CD4^+^ T cells (Fig. 2g-i, Table S3), but importantly not with the signatures of T cell activation (Fig. 2h), again indicating that the absence of Treg suppression did not result in wholesale T cell activation. Indeed, a Treg activation signature ^51^ was downregulated in IPEX cells (Fig. 2g). Intriguingly, transcripts differentially expressed in tumor-infiltrating Tregs (2 independent studies) were counter-regulated in IPEX CD4+ T cells, as were interferon-stimulated genes (Fig. 2g,i). This broad change was consistent with the up-regulation of major response regulators like *RELB*, *KRAS* or *HIF1*. Overall, our results indicated that the Treg signature was in large part maintained in Tregs from IPEX patients, albeit with a notably high degree of variability. More unexpected was the peculiar transcriptomic footprint shared by IPEX Tregs and Tconvs, which might result from integration of extracellular cues, but not from a generic T cell activation state.

### Different types of Tregs in IPEX patients

From these preliminary indications, we proceeded to the crux of the work, applying scRNAseq to identify with unbiased profiling the actual types of FOXP3-deficient Treg-like cells, which might be blurred by the averaging inherent in population profiling, or have escaped recognition because of shifts in the CD25/CD127 markers. We sorted total CD4^+^ T cells, to yield granular information on both Treg and Tconv pools, and performed scRNAseq on the two cohorts as above (a total of 52,776 cells passing QC (Table S4) from 11 IPEX and 11 HD donors). All results were observed in both the initial and replication cohorts, but are combined below for simplicity. Experimental confounders were minimized by multiplexing IPEX and HD samples in the same scRNAseq runs ^52^. Residual inter-cohort and inter-experiment effects were corrected using the Canonical Correlation Analysis (CCA) and k-nearest neighbor-based integration methodology ^53^ (Fig. 3a).

**Fig. 3.**
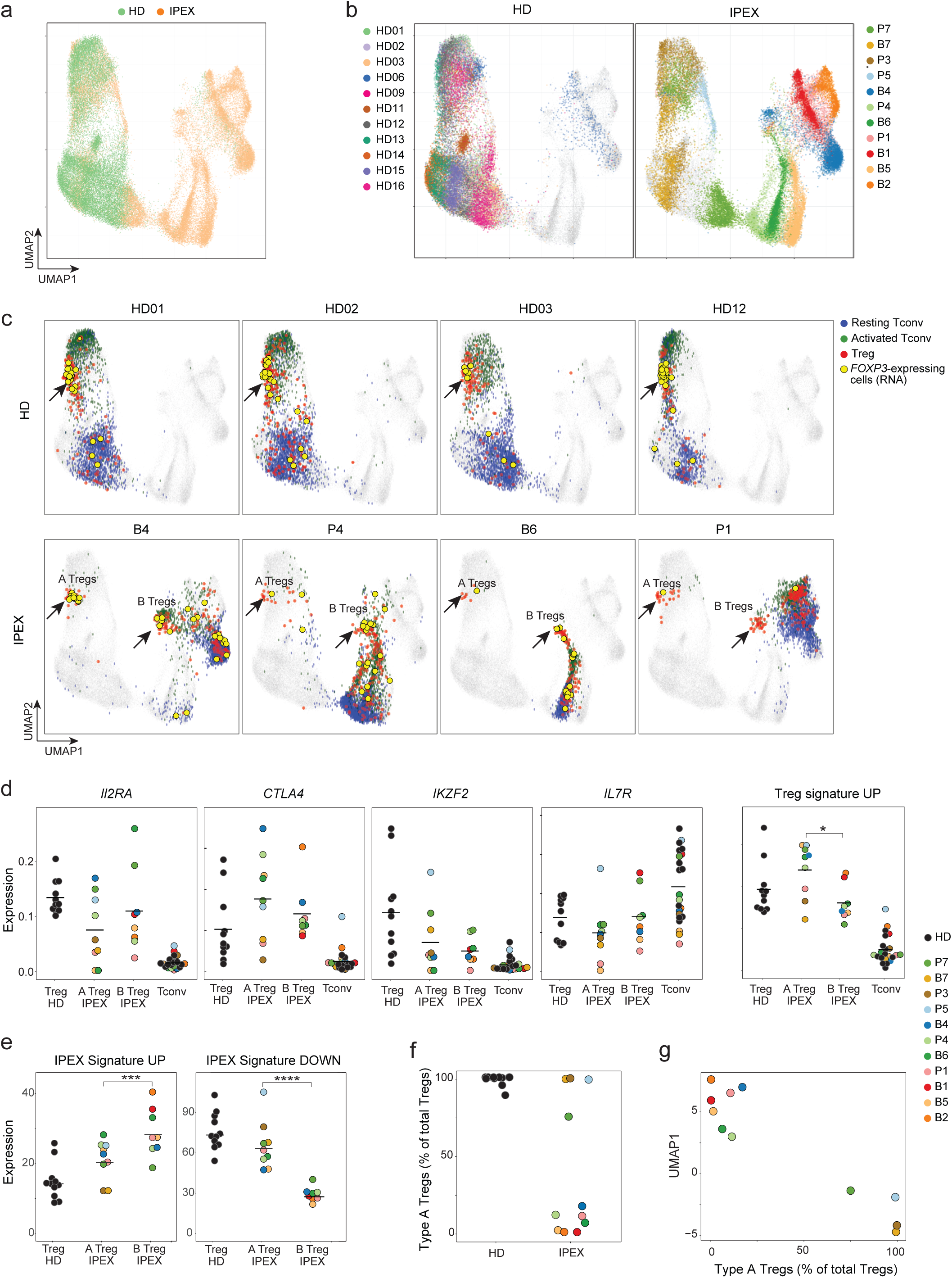
scRNAseq reveals the heterogeneous effect of FOXP3 deficiency in IPEX Tregs. **a.** scRNAseq was performed on CD4+ T cells, and results displayed as a 2D UMAP plot of all CD4^+^ cells from IPEX and HD samples (52,776 cells altogether, merged with Canonical Correlation Analysis and knn-graph, see Methods) **b.** Same UMAP as a, color-coded by individual donor. **c.** Same UMAP as a, with four representative HD and IPEX donors (see Fig. S8 for other donors). Blue, green, and red cells are resting Tconvs, activated Tconvs, and Tregs, respectively. *FOXP3*-expressing cells (RNA) are yellow. Type A and B Tregs in IPEX donors are indicated by an arrow. **d.** Normalized counts expression of *IL2RA*, *CTLA4*, *IKZF2*, *IL7R,* and the Treg signature in Tregs from HD and IPEX (type A and B) and Tconv cells; each dot is a sample. **e.** Average expression of the IPEX signature in type A and B IPEX Tregs and HD Tregs (normalized counts). **f.** Proportion of type A Tregs in total Tregs for each sample. **g.** Proportion of Type A Tregs plotted against the average UMAP1 dimension for each sample.

In the UMAP projection of the integrated data, CD4^+^ T cells partitioned sharply according to disease: cells from HD donors clustered together, while those from IPEX patients were much more dispersed (Fig. 3a), each tending to form an island distant to various degrees from the HD group (Fig. 3b). These distances were not batch artefacts (similar HD/IPEX partitions were seen in each batch, Fig. S6a). Replicate samples from 3 patients drawn >1 year apart mapped to the same regions, as did samples from patient P7 collected before and after immunosuppressive treatment (Fig. S6b), indicating that the different locations were patient-specific and not consequences of environmental or treatment variables. Conventional and scRNAseq reflected the same perturbation, as UMAP1 scores axis that partitioned IPEX and HD samples corresponded with the IPEX signature above (Fig. S6c).

We then sought to deconvolute, at single-cell resolution, Tregs among these CD4^+^ T cells. Tregs were identified in HD and IPEX samples in an unsupervised approach, using community detection in the reciprocal PCA integration network (implemented in Seurat-V3 ^53^), also supported by activity at the *FOXP3* locus. Three distinct populations of CD4^+^ T cells could be identified (S6d): two Tconv populations (resting and activated) and one Treg (confirmed by expression of Treg signature genes *FOXP3, IKZF2, CTLA4, IL2RA, TIGIT*; Fig. S7a). When Tregs thus identified were displayed onto the UMAP space, HD Tregs formed a tight cluster for all HD samples (Fig. 3c), which co-localized with *FOXP3* expression (yellow dots on Fig. 3c). In IPEX samples, Treg-like cells were similarly identified, in proportions equivalent to controls, even for fully FOXP3-deficient P1 (Fig. S7b), but they split into two groups (Fig. 3c, S8a): some (hereafter referred to as type-A IPEX Tregs) mapped in the same cluster as HD Tregs, while others clustered away, in their respective patient-specific “island” (type-B). Both were truly Tregs, expressing *FOXP3* mRNA (Fig. 3c), core Treg transcripts (*IL2RA, CTLA4, IKZF2,* low *IL7R*) and the Treg signature (Fig 3d). Type-A and type-B Tregs did differ, with higher levels of Treg signature genes in type-A Tregs, and higher representation in type-B of the IPEX signature (Fig. 3e, Fig. S8b). Finally, proportions varied between IPEX patients (Fig. 3f): type-A Treg dominated in some, almost to HD levels, but were far less abundant in others. This proportion was stable in independent samplings of the same patient, and related to the intensity of the IPEX signature in Treg and Tconv cells: patients with the highest index had the lowest fraction of type-A Tregs (Fig. 3g).

Thus, scRNAseq analysis revealed a subset of Tregs that closely resemble normal Tregs, and another with more extensive perturbations. The origin of these two distinct Treg populations in IPEX patients was unclear. To exclude maternal microchimerism (wild-type Treg cells from the mother could have a competitive advantage in the IPEX offspring), we checked female-specific *XIST* transcripts: no *XIST*-positive cells were found among any patient’s Tregs. A striking feature of type-A Tregs was their marked downregulation of the IPEX signature.

### IPEX does not affect Tconv phenotypes

Tregs can affect Tconv polarization in many ways ^15^, and it is generally assumed that Treg deficiency in IPEX patients leads to excessive Tconv activation and differentiation, and an over-representation of their effector states ^54, 55^. Unsupervised clustering of Tconv cells performed in the shared reciprocal PCA space, after regressing out the IPEX effect, distinguished 9 clusters (5 resting and 4 activated/memory types, judging from the expression of characteristic markers and transcription factors like *CCR7, SATB1, CD69, TBX21, GATA3*) (Fig. 4a). Their distribution, and the expression of defining transcripts, was strikingly similar for HD and IPEX samples (Fig. 4a,b). This observation was confirmed by flow cytometry, which showed similar ranges of CD45RA^+^ cells among CD4^+^ T cells in IPEX and HD PBMCs (Fig. 4c). Thus, beyond the shared signature, IPEX disease and immunosuppressive treatments seemed to have limited impact on other phenotypic aspects of circulating Tconv cells.

**Fig. 4.**
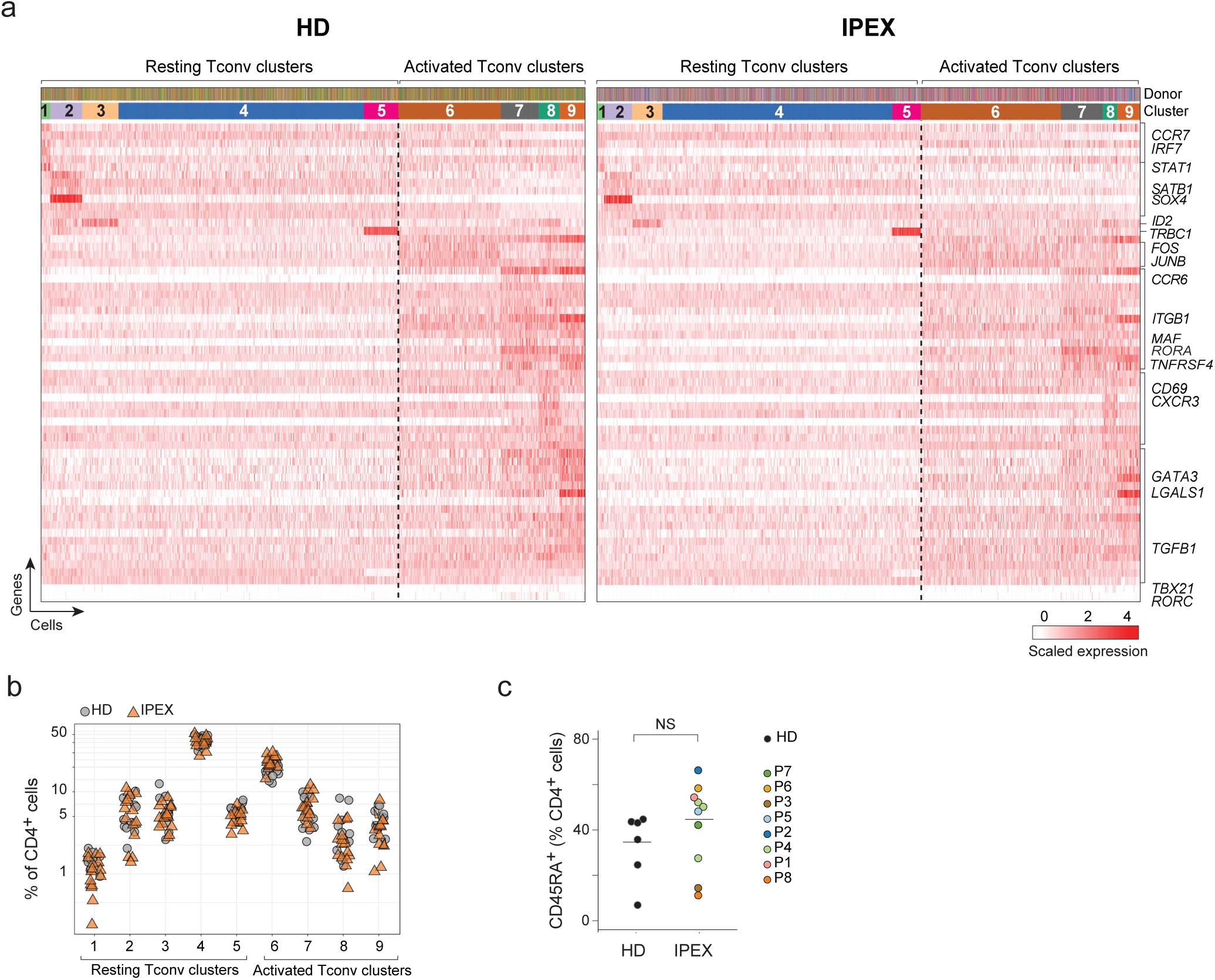
Normal Tconv phenotypes in IPEX patients. **a.** From the scRNAseq Tconv datasets (HD and IPEX combined), the IPEX signature was regressed out (see Methods) and biclustering was performed to define clusters in resting and activated Tconv. plotted as a heatmap for expression of the most characteristic genes for each Tconv cluster (each vertical line represents one cell). Top ribbons indicate donor origin and cluster annotations for every cell. **b.** Proportion of resting and activated Tconv clusters in total CD4+ cells in HD and IPEX in scRNAseq data. **c.** Proportion of CD45RA+ (resting) CD4+ T cells determined by flow cytometry in IPEX and HD.

### Mixed populations of *Foxp3*-active cells in *Foxp3*-deficient mice

The *FOXP3* deficiency in IPEX patients thus led to a global change affecting all CD4+ T cells, and to a diversity of Treg-like cells. To assess mutational impacts in a setting devoid of genetic or therapeutic variables, we performed parallel analyses on *Foxp3*-deficient mice [the previously described *Foxp3^ΔEGFPcre^* inactivating reporter line ^41^, hereafter denoted “Δ*Foxp3*”], in which FOXP3 protein is absent, but the active *Foxp3* locus can be detected via the reporter.

We used scRNAseq to analyze the diversity of CD4^+^ T splenocytes in Δ*Foxp3* mice and WT controls (4 mice per group, 18,000 cells altogether, again multiplexing samples in the same runs, Table S4). As in human patients, cells from WT and Δ*Foxp3* mice clustered separately on the UMAP projection, here again reflecting the generic ΔFoxp3 signature (Fig. 5a) present in every mutant mouse (Fig. S9a). Unsupervised clustering in the shared CCA space identified, for WT CD4^+^ cells, resting and activated Tconv and a tight group of Treg cells (blue, green and red, respectively, Fig. 5b top). These assignments were confirmed by the location of sorted Tconv or Treg cells (Fig. 5b, top right), by differential expression of canonical genes (e.g. *Ccr7, Sell, Cd44, Tbx21*; Fig. S9b), and at the *Foxp3* locus (as *Cre* and *Foxp3* transcripts; Fig. 5b, bottom). Note how Tconv clusters were similarly structured in Δ*Foxp3* and WT mice, as they had been in human patients (Fig. S9b).

**Fig. 5.**
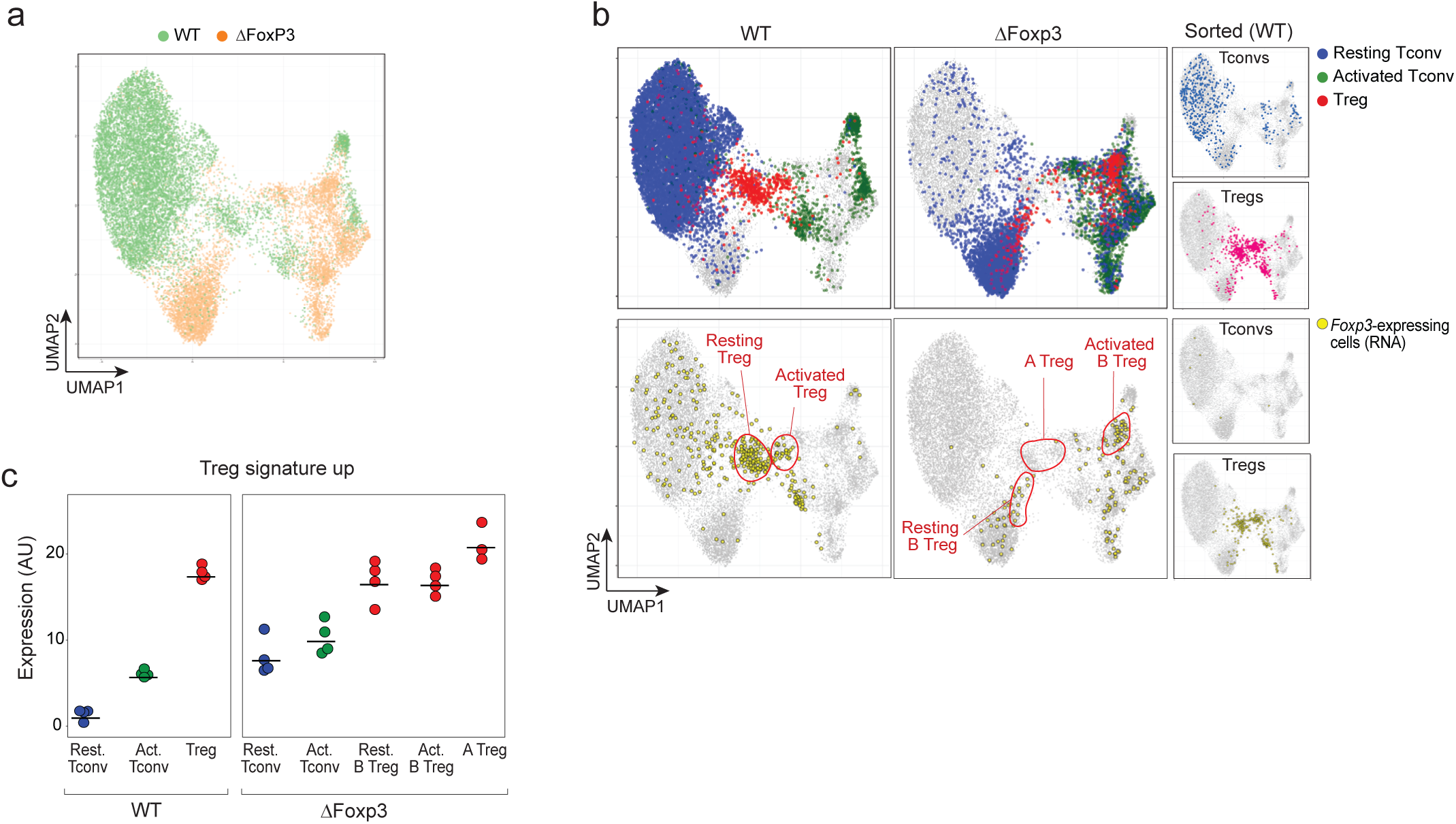
scRNAseq reveals heterogeneous effects of the *Foxp3* ablation in Δ*Foxp3* mice. scRNAseq was performed on CD4+ T cells from WT and Δ*Foxp3* mice (18,097 cells altogether). **a.** 2D UMAP plot of CD4+ single-cell transcriptomes for WT (green) and ΔFoxp3 (orange) (n = 4 mice per group, 2 experiments). **b.** Same UMAP as b. Top row: resting Tconv (blue), activated Tconvs (green), and Tregs (red) (assigned by graph-based clustering on the merged dataset after regressing out the Δ*Foxp3* signature, see Methods) are highlighted; bottom: cells with an active *Foxp3* locus (*GFP* or *Cre* transcripts detected) are colored yellow. The small inserts on the right show sorted control WT Tconvs and WT Tregs, sorted and included as spike-in controls. **c.** Treg signature expression in resting Tconvs, activated Tconvs, and Tregs in WT and Δ*Foxp3* mice (normalized counts).

As for IPEX patients, Treg-like cells of Δ*Foxp3* mice were multiform (Fig. 5b): a minor fraction of “A-Tregs” closely resembled normal Tregs, in the same small proportions as in the most complete IPEX deficiencies; a larger proportion of “B-Tregs” that mapped into resting and active areas of the UMAP projection (Fig. 5b), also expressing *Foxp3* (Fig. 5b, bottom). Both expressed the Treg signature, highest for Type-A Tregs (Fig. 5c). Together, these Treg-like populations accounted for 7% of total T CD4^+^ cells, as in WT Tregs (Fig. S9c). Thus, Treg-like cells in *Foxp3*-deficient mice showed the same heterogeneity as in human IPEX patients: a mix of healthy and altered Tregs, but with a dominant transcriptional signature that cut across both Treg and Tconv CD4+ T cells.

### Cell-intrinsic and extrinsic impact of *Foxp3* deficiency

In humans and mice, FOXP3 deficiency resulted in a heterogenous mix of Treg-like cells, and a strong disease-specific signature unexpectedly shared by Tregs and Tconvs. The latter suggested cell-extrinsic influences on the transcriptomes of CD4^+^ T cells, which we investigated by analyzing bone marrow chimeras (BMC) constructed by transfer of equal proportions of congenically-marked stem cells from WT and Δ*Foxp3* mice, a protocol that prevents the *scurfy*-like disease that appears after reconstitution with *Foxp3*-deficient stem cells ^24^. Cell-extrinsic transcriptional hallmarks should be reverted in genetically *Foxp3*-deficient cells by the presence of WT cells in the same mouse. Ten weeks after reconstitution, 5,556 CD4^+^ T cells from either Δ*Foxp3* or WT compartments of BMC mice were profiled by multiplexed scRNAseq (3 mice/group, data aligned to the same UMAP space as above).

In this mixed setting, many of the characteristics of Δ*Foxp3* CD4+ T cells essentially disappeared (Fig. 6a), while WT cells were unchanged. First, on the UMAP plot, resting and activated Tconvs from Δ*Foxp3* compartments shifted and overlapped with WT cells from the same mice (Fig. 6a). Accordingly, the Δ*Foxp3* signature was flattened out for mutant cells in the BMC setting (Fig. 6b, S10a). These results demonstrated the cell extrinsic origin of the Δ*Foxp3* signature in *Foxp3*-deficient Tconv.

**Fig. 6.**
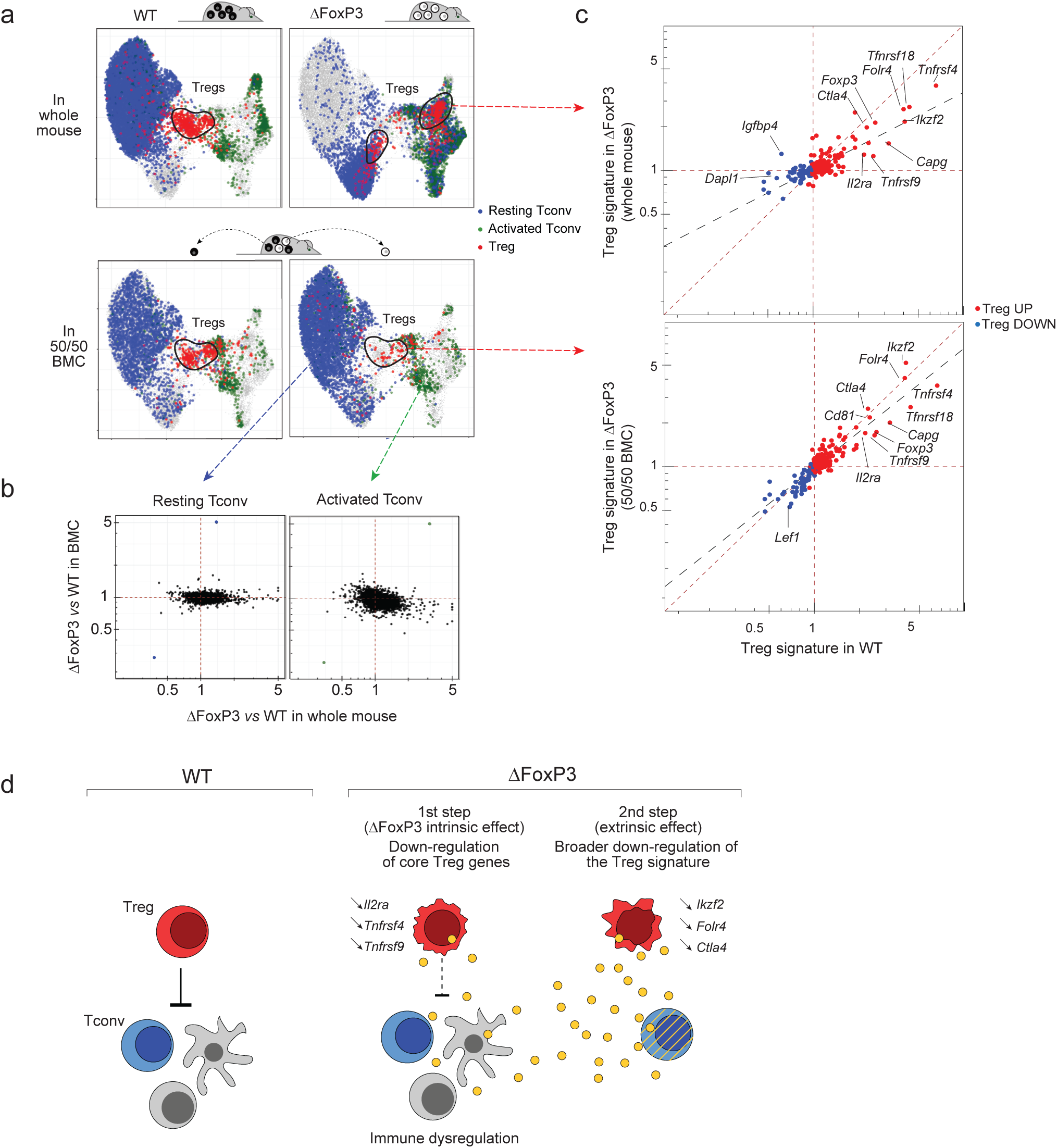
Cell intrinsic and extrinsic effect of *Foxp3* deficiency in Tregs and Tconvs. scRNAseq was performed on sorted and hashtagged CD4+ T cells from 50/50 mixed BMC mice (WT and Δ*Foxp3* hematopoietic stem cells, congenically marked) (8,556 cells altogether) **a.** Same UMAP as 5b. integrating WT and Δ*Foxp3* CD4+ cells from whole mice (top) or from chimeras (bottom row). Resting Tconvs (blue), activated Tconvs (green), and Tregs (red) (identified by graph-based clustering on the merged dataset after regressing out the Δ*Foxp3* signature, see Methods) are color-coded as in Fig. 5. **b.** scRNAseq data for Tconv cells were collapsed, and expression ratio between WT and Δ*Foxp3* Tconv calculated for whole mice (x-axis) or for 50/50 BMC (y-axis); for the latter, WT and Δ*Foxp3* cells originated from the same host. **c.** Treg/Tconv ratio in Δ*Foxp3* cells (y-axis) in whole mice (top) and in 50/50 BMC (bottom), both plotted against the same Treg/Tconv expression ratio in WT (x-axis). Treg signature genes are highlighted. Dashed lines (linear regression) represent the slope of the fit. **d.** Two-step model of the impact of FOXP3 deficiency in mice and humans. First, the absence of FOXP3 downregulates a few core Treg signature genes in Tregs (cell-intrinsic effect), leading to perturbed control of other immune cells (which might include known Treg targets like Tconv, DCs, NKs, or other), unleashing a secondary inter-cellular response (e.g. cytokines, cell surface modulators) which imprints the dominant IPEX signature on Treg and Tconv, and further affects Treg functionality, amplifying disease pathology.

As expected, Δ*Foxp3* cells were partially outcompeted by WT Tregs (1:3) in the BMC mice (Fig. S10b), but their phenotypes were also corrected: they were all type-A Tregs that overlapped with WT Tregs on the UMAP (Fig. 6a), and their expression of the Treg signature was almost completely normalized (Fig. 6c). Only 13 genes remained under-expressed in the Δ*Foxp3* Treg compartment, including classic FOXP3 target loci (*Il2ra, Tnfrsf4, Tnfrsf9, Tnfrsf18, Capg*); no genes were upregulated, confirming that FOXP3 is mainly a transcriptional activator. The other Treg signature genes were otherwise normally expressed, contrasting with their general downregulation in Δ*Foxp3* mouse.

In conclusion, the dominant suppressive effect of WT Tregs curtailed the Δ*Foxp3* signature in Treg and Tconv, revealing a narrower intrinsic effect of *Foxp3* on a minor proportion of the Treg signature, highlighting both intrinsic and extrinsic consequences of the *Foxp3* deficiency in the disease context. We thus propose a “two-step” model of the IPEX/*scurfy* disease (Fig. 6d). First, the intrinsic effect of *Foxp3* in Tregs dysregulates a few core Treg genes, which unleashes a systemic response. Second, this milieu imprints broad changes on both Treg and Tconv (IPEX/ΔFoxp3 signatures), and further destabilizes Treg signature expression and dampening Treg effector function.

## DISCUSSION

These results provide unprecedented cellular and genomic resolution of a primary immunodeficiency, and reveal unexpectedly multifarious molecular and cellular consequences of FOXP3 deficiency. There were surprisingly limited cell-intrinsic perturbations in Treg cells, associated with a dominant and monomorphic signature of cell-extrinsic origin, that cuts across the transcriptome of all CD4^+^ T cells and amplifies Treg perturbations.

We thus propose a two-step model of the disease’s molecular pathogenesis. FOXP3 is actually important for only very few Treg genes: when the milieu was normalized by the presence of WT Tregs the cell-intrinsic impact of the *ΔFoxp3* deficiency was exerted on only a handful of genes (*Il2ra*, *Tnfrsf4*, *Tnfrsf9*, *Tnfrsf18*, *Capg*). This surprisingly short list corresponds to a “core set” of genes that are expressed by all Tregs ^7^, are directly transactivated by and bind FOXP3 ^14, 56, 57^. These genes encode the major homeostatic regulator of Treg cells (*IL2RA*), and several members of the TNFR superfamily, which are also connected to Treg homeostasis and function ^58, 59^. We propose that their downregulation is the first step (accordingly, *IL2RA* deficiency also causes an IPEX-like disease). This initiates a systemic reaction which constitutes the second step of the FOXP3 deficiency syndrome: a broadly shared signature that marks both Treg and Tconv cells, further perturbs Treg signature transcripts, and amplifies in a vicious circle the defect in Treg suppressive activity.

What, then, is this IPEX signature, the hundreds of genes that were impacted equivalently in Treg and Tconv cells? It was shared among all IPEX patients, with different intensities rather than qualitative differences, stressing its common etiology. It proved stable over several years in each multiply-tested patient, before and after (and unrelated to) treatment. The genes involved were not simply T cell activation genes, as might have been expected from a loss of suppression (activation-induced transcripts were actually repressed). We hypothesize that it is due to an inductive element(s), delivered via cytokine or cell-cell contact, normally repressed by Tregs but unleashed by the FOXP3 deficiency. This signal might originate from other T cells, or from other immunocytes, much as dendritic and NK cells are the first responders to acute Treg ablation in DTR systems ^60–62^. The presence of WT Tregs in mixed BMC mice would restore the negative feedback, thus reinstating better Treg function, evoking the “infectious tolerance” concept ^63^, the presence of normal suppressors improving the tone and function of defective ones. Pathway and ontology analysis revealed no clear match, except for effects on a few IFN- and TNF-induced genes. There was, on the other hand, an intriguing anti-correlation for genes over-expressed in tumor-infiltrating Tregs ^64, 65^), CD4+ T cells from IPEX patients turned off much of the tumor Treg signature.

This principal effect cutting across both Treg and Tconv evokes the debated hypothesis that FOXP3 deficiency has an intrinsic effect in Tconv cells, associated with the transient induction of FOXP3 in activated Tconv ^45–48^. The mixed chimeras demonstrated that the FOXP3 deficiency affected Tconv cell-extrinsically. However, it remains an open question whether this disease signature contributes to pathology via further dampening of Treg function, or through Tconv defects.

That FOXP3 is not absolutely required for Treg differentiation and homeostasis is now well established, with the existence of Treg-like cells described repeatedly in mice and humans lacking FOXP3 ^39–44^. These “Treg wannabes” were reported to maintain some Treg features (self-reactivity, partial Treg signature, activity at the *FOXP3* locus) while acquiring some Tconv characteristics (no *in vitro* anergy, cytokine expression). The unexpected insight emerging from our single-cell analysis of deficient humans and mice was the wide array of cells with Treg-like characteristics and/or an active *FOXP3* locus. Rather than a single population of Treg wannabes, several distinct populations were present: first, a large component of B-Tregs, with many Treg transcriptional characteristics, but perturbed by the IPEX signature. Second, and most mysterious, the subset of A-Tregs that seemed almost unaffected relative to WT Tregs: full Treg signature, active *FOXP3*, and no IPEX signature, as if these cells had somehow become non-responsive to the systemic influence, generating “escape variants”, e.g. by dampening receptors or signaling, or because they reside in protected niches. But, then, why only for Tregs?

These results show that it can be misleading to infer the transcriptional footprint of a TF, and its mechanistic causality in disease, solely from the transcriptome of deficient cells, since it can be perturbed as here by dominant cell-extrinsic effects. When the extrinsic effects were blocked in the chimeras, the true core signature of *Foxp3* proved much smaller than the first analysis of the deficiency had suggested. IPEX is a rare disease, and our cohorts were not powered to robustly detect clinical correlates of the patients’ cellular and transcriptional characteristics. On the other hand, there was no obvious correlation between clinical severity indicators and the integrated gene expression metrics. This finding is congruent with the notion that molecular severity of the FOXP3 mutations in IPEX patients only loosely correlates with clinical severity ^34, 66^ - here, the null mutation in patient P1 did not determine the most severe disease.

There are some implications of our findings for therapeutic strategies in IPEX. Current management, when bone marrow transplantation is not an option, involves immunologic dampening via immunosuppressants. Our results might suggest harnessing those type A Tregs that are present in the patients by sustaining their homeostasis, or complementing the small set of primary FOXP3 targets identified here (e.g. with IL2 or muteins thereof, or with TNFR family agonists). Alternatively, damage might be avoided by blocking the signaling mechanism that imparts the dominant IPEX signature to all T cells.

In conclusion, the new landscape of Treg and Tconv cells revealed in IPEX patients by single-cell analysis and their correction in BMC, have opened a new perspective on the disease, and on the role of FOXP3 and Tregs in immune homeostasis.

## ACKNOWLEDGEMENTS

We thank: Drs. Megan Levings and Sasha Rudensky for insightful discussions; K. Hattori, C. Araneo, K. Seddu, and the Klarman Cell Observatory team, for help with mice, cell sorting and single-cell profiling. This work was funded by grants from the NIH to CB&DM (AI116834, AI125603), TAC (AI085090), the Institut National de la Santé et Recherche Médicale, the European Union Seventh Framework (#269037 and #261387) and Horizon 2020 (#693762), the Agence Nationale pour la Recherche (Investissement d’Avenir ANR-10-IAHU-01) to IA, ES, MD, JL, MC, BN, FRL, FR and NCB. JL was supported by by an INSERM Poste d’Accueil and an Arthur Sachs scholarship.

This paper has not been peer reviewed.

## AUTHOR CONTRIBUTIONS

DZ and LMC performed the experiments; ES, SK, MD, SB, JZ, KC, BN, MIGL, FR, MCB, FRL, M, IA, TAL, LMC provided samples and discussed interpretations; DZ, LMC, TAL, IA, CB and DM designed the study, analyzed and interpreted the data; DZ, JL, CB and DM wrote the manuscript.

## SUPPLEMENTARY METHODS

### Mice

*Foxp3^ires-gfp^*/B6 ^67^ and *Foxp3^ΔEGFPiCre^xR26^-YFP^* (Δ*Foxp3*) mice were maintained in our colony. Except when specified, 3 week-old male mice were used in this study (Table S4). Mice were housed under specific pathogen-free conditions and all experimentation was performed following the animal protocol guidelines of Harvard Medical School and Boston Children’s Hospital (HMS IACUC protocol 02954).

#### Bone marrow chimeras

C57BL/6J mice were irradiated with 10 Gy and reconstituted with equal proportions of congenically labeled T cell-depleted bone marrow cells from WT (*Foxp3^ires-gfp^*x*CD45.1*) and Δ*Foxp3* (*Foxp3^ΔEGFPiCre^xR26^-YFP^xCD45.1/CD45.2*) mice. Bone marrow cells were harvested from femurs, tibias and hip bones from two WT and two Δ*Foxp3* male mice. After red blood cell lysis with ACK for 1 min at 4°C, T cells were depleted: bone marrow single-cell suspensions were incubated with 20 uL of biotinylated anti-CD3e antibodies (OKT3, Biolegend) for 10 min in 2 mL of MACS buffer (PBS, FBS 0.5%, EDTA 2mM), washed, and then incubated with 200 μL of streptavidin beads (Dynabeads Biotin Binder, 11047; Thermo Fisher Scientific) for 20 min in 5 mL of MACS buffer. Isolation of the CD3-negative population was performed after three magnet incubations for 2 min. A total of 4 million cells (2 million WT and 2 million Δ*Foxp3*) were injected i.v. in each mouse. Mice were treated for 2 weeks with trimethoprim-sulfamethoxazol and analyzed 10 weeks later.

### Human cohorts

Male IPEX and healthy donor whole-blood samples were obtained under protocols reviewed and approved by the local Institutional Review Boards at each center (Boston Children’s Hospital 04-09-113R, Necker/Imagine C 15-13_CODECOH_AR, HMS IRB15-0504). Anonymized clinical data included age at onset and at blood sampling, clinical symptoms (enteropathy, diabetes, eczema, other autoimmune diseases and allergy), ancillary testing, treatments and long-term outcomes (Table S1). Two independent cohorts of samples were profiled (Table S1). Each cohort contains samples from both centers. Cohort 1 samples were processed in two different scRNAseq runs, cohort 2 (replication cohort) in three scRNAseq runs. Samples from each cohort were sorted and sequenced as one batch for population RNA sequencing (Table S4).

### Peripheral blood mononuclear cells (PBMC) isolation

Whole blood was collected in K_2_ EDTA tubes and processed within a few hours. An equal volume of room-temperature PBS/2mM EDTA was mixed into 15ml of blood and carefully layered over 14ml Ficoll Hypaque solution (GE Healthcare). After centrifugation for 30 min at 900 g (with no break), at room temperature, the mononuclear cell layer was washed three times with excess HBSS (Gibco) (10min at 400 g) and resuspended in 2ml of HBSS. The pellet was resuspended in 90% FBS-10% DMSO, 20 million cells/mL, 1mL per vial, cooled progressively in isopropyl alcohol (Mr. Frosty™ Freezing Container, Thermo Fisher) for 24h and stored in liquid nitrogen for long term.;

Vials were thawed in 10 mL 10% FBS RPMI, cells washed (500g for 5 min) and resuspended in FACS buffer (phenol red–free RPMI, 2% FBS, 0.1% azide and 10mM HEPES, pH 7.9). After cell counting, the samples were allocated for flow cytometry, population RNA sequencing and scRNAseq.

### Flow cytometric profiling of IPEX and HD

Cells were stained for flow cytometry in 100 µL of FACS buffer (phenol red–free DMEM, 2% FBS, 0.1% azide and 10mM HEPES, pH 7.9) for 10 min with 10uL FcBlock (supernatant of clone 2.4G2, ATCC HB-197, hybridoma cultures) and the following antibodies: CD3 BV605 or A700 (2µL, OKT3, Biolegend), CD4 PerCP-Cy5.5 (2 µL, OKT4, Biolegend), CD25 PE-Cy7 (3 µL, BC96, Biolegend), CD127 A488 (3 µL, A019D5, Biolegend), CD45RA PB (3µL, HI100, Biolegend). After permeabilization/fixation for 2 h on ice, FOXP3 (APC, clone PCH101; Biolegend; 2 µL) and anti-HELIOS PE (PE, clone 22F6, Biolegend, 2 µL) staining was performed overnight at 4°C in the dark in accordance with the manufacturer’s instructions (eBioscience™ FOXP3 / Transcription Factor Staining Buffer Set). Data was recorded on an LSRII flow cytometer (BD Biosciences) and analyzed using FlowJo v10. FOXP3 MFI was normalized to the mean of healthy donors. To generate tSNE plots from the flow cytometry data, compensated, scaled data in the Lymphocyte/Singlet/CD3^+^/CD4^+^ gate were exported in .csv format from FlowJo. In R, the matrix containing CD3, CD4, FOXP3, HELIOS, CD25, CD127, CD45RA expression was centered and scaled (by marker) before performing PCA using the *prcomp* function, and tSNE projections calculated with Rtsne(dims = 2, perplexity = 50, check_duplicates = F, pca = F, max_iter = 500).

### Low-input RNA sequencing of human and mouse samples

*RNA-seq* was performed with the standard ImmGen low-input protocol (www.immgen.org). Human Tregs and Tconvs were sorted as DAPI^−^ CD4^+^CD25^hi^CD127^lo^ and CD4^+^CD25^-^CD127^hi^, respectively on a Moflo Astrios Cell Sorter (Beckman Coulter). Mouse Tregs and Tconvs were sorted as DAPI^−^TCRβ^+^CD4^+^G/YFP^+^ DAPI^−^TCR^+^CD4^+^G/YFP^-^, respectively. A total of 1,000 cells were double-sorted directly into 5ul of lysis buffer (TCL Buffer (QIAGEN) supplemented with 1% 2-Mercaptoethanol). Smart-seq2 libraries were prepared as previously described ^68^ with slight modifications. Briefly, total RNA was captured and purified on RNAClean XP beads (Beckman Coulter). Polyadenylated mRNA was then selected using an anchored oligo(dT) primer (50 – AAGCAGTGGTATCAACGCAGAGTACT30VN-30) and converted to cDNA via reverse transcription. First strand cDNA was subjected to limited PCR amplification followed by Tn5 transposon-based fragmentation using the Nextera XT DNA Library Preparation Kit (Illumina). Samples were then PCR amplified for 12 cycles using barcoded primers such that each sample carries a specific combination of eight base Illumina P5 and P7 barcodes for subsequent pooling and sequencing. Paired-end sequencing was performed on an Illumina NextSeq 500 using 2 x 38bp reads with no further trimming.

Reads were aligned to the human genome (GENCODE GRCh38 primary assembly and gene annotations v27) with STAR 2.5.4a (https://github.com/alexdobin/STAR/releases). The ribosomal RNA gene annotations were removed from GTF (General Transfer Format) file. The gene-level quantification was calculated by featureCounts (http://subread.sourceforge.net/). Raw read counts tables were normalized by median of ratios method with DESeq2 package from Bioconductor (https://bioconductor.org/packages/release/bioc/html/DESeq2.html) and then converted to GCT and CLS format. Samples with less than 1 million uniquely mapped reads were excluded from normalization to mitigate the effect of poor quality samples on normalized counts. Genes with a minimum read count of 5 in all replicates of a population (31,448 human genes) were retained. A pseudo count of 1 was added and log2-transformed prior to quantile normalization. Quantile-normalized counts were converted back to a linear scale.

*IPEX signature* genes were identified by computing the ratio of expression in Tregs or Tconv of all IPEX patients vs all healthy donors (p.value as simple uncorrected t.test). *IPEX-up indices* for each individual were calculated by selecting the 100 transcripts most over-expressed in IPEX Tregs overall (FC>3), calculating their differential expression in Treg cells of each patient relative to the mean of all HD Tregs, and averaging the log2 of these foldchanges (and similarly for IPEX-down indices from transcripts with IPEX/HD <0.37). The TregUp indices were similarly calculated (expression in Tregs of each individual over mean expression in all healthy donors, average the log2 of these foldchanges).

#### Geneset enrichment analysis

Gene signatures were curated from published datasets (references in the signature name) ^7^. The Human and Mouse Treg signatures have been reported ^5, 6^. Data were downloaded from the Gene Expression Omnnibus (GEO) (https://www.ncbi.nlm.nih.gov/geo/). Only datasets containing replicates were used. To reduce noise, genes with a coefficient of variation between biological replicates <0.6 in either comparison groups were selected. Up- and downregulated transcripts were defined as having a fold change in gene expression >1.5 or <0.6 and a t-test *p*-value <0.05. Other signatures were obtained from MSigDB C7 Immunologic signatures collection ^69^. Geneset enrichment analysis with the IPEX signature was performed using the hypergeometric distribution and type I error was controlled using FDR. Signatures with *p*-value <0.001 (all with FDR <6%) are reported.

### scRNAseq analysis of human PBMC samples

scRNAseq was performed in several batches (different experiment dates in Table S1 and S4): two for cohort 1 and three for cohort 2. Cohort 1 samples were profiled with the 10X Genomics Single Cell 3′ Reagent Kit (V2 chemistry), Cohort 2 samples with the 10X Genomics Single Cell 3′ Reagent Kit (V3 chemistry) and sample barcoding with DNA-tagged antibodies (“hashtagging”) ^52^. See also Table S4.

#### Cell sorting and pooling using cell hashtagging

Cells were stained in 100 µL of FACS buffer (phenol red–free RPMI, 2% FBS, 0.1% azide and 10mM HEPES, pH 7.9) for 10 min with 10uL FcBlock (homemade) and the following antibodies: CD3 A700 (2µL, OKT3, Biolegend), CD4 PerCP-Cy5.5 (2 µL, OKT4, Biolegend), CD25 PE-Cy7 (3 µL, BC96, Biolegend), CD127 A488 (3 µL, A019D5, Biolegend), 5 uL (2.5ug) of a unique hashtag antibody (TotalSeq™-A0251 to A0258 anti-human Hashtag 1 to 8 Antibody). 8,000 DAPI^−^CD3^+^CD4^+^ T cells were single sorted using the Moflo Astrios Cell Sorter (70 um nozzle, Beckman Coulter) in 30 ul of PBS-BSA 0.1%. Samples with different hashtag antibodies were sorted in the same tube and the total volume was adjusted to 30 uL.

#### scRNAseq libraries (10x Genomics)

Cells were encapsulated in one channel per sample (cohort 1 runs) or in 1 channel per pool (cohort 2 samples) of a 10x Chromium instrument, and libraries were constructed with the Single Cell 3′ Reagent Kit (V2 for cohort 1 and V3 chemistry for cohort 2) (https://support.10xgenomics.com/single-cell-gene-expression/libraryprep/). Libraries were sequenced on the NextSeq 500 platform (28/8/0/91, Read1/i7/i5/Read2). Gene counts were obtained by aligning reads to the hg38 transcriptome using CellRanger software (v3.0.2) (10X Genomics) (default parameters).

#### Hashtag libraries

Hashtag libraries were made separately as described in Stoeckius et al. ^52^ (https://citeseq.files.wordpress.com/2019/02/cell_hashing_protocol_190213.pdf). In brief, at the cDNA amplification step in the Single Cell 3′ Reagent Kit protocol, the yield of HTO products was increased using an “additive” primer to cDNA PCR. Hashtag-derived cDNAs (<180bp) and mRNA-derived cDNAs (>300bp) were then separated using 0.6x SPRI bead selection. The supernatant contains the Hashtag-derived cDNA that is purified with two rounds of 2x SPRI beads. The sequencing oligos are added by PCR which also amplifies the Hashtag library. Libraries were sequenced on the NextSeq 500 platform (28/8/0/91, Read1/i7/i5/Read2). Hashtag count matrices were obtained from CITE-Seq-Count package (https://zenodo.org/record/2590196) (default parameters). Each droplet from the Gene Count matrix was matched to a Hashtag using the HTODemux function from the Seurat v3.1.2 package. Doublets (droplets with 2 hashtags) were excluded and cells were assigned to the max hashtag signal. The Hashtag count data was also analyzed by tSNE for a visual check (clear separated clusters for each hashtag). All single cells from the gene count matrix were matched unambiguously to a single hashtag (and therefore their original donor).

#### Quality control

Cells with less than 500 (V2) or 1000 (V3) counts (V2) were excluded. Dead cells with more than 10% of counts mapping to mitochondrial genes were excluded. Doublets were excluded using scrublet (doublet score > 0.3) ^70^. Finally, cells were annotated using singleR ^71^ with the BluePrintEncode reference data. Cells that were not annotated as CD4+ cells were excluded.

#### Batch correction

Batch correction was performed using the Integration method in Seurat V3 as described in Stuart et al. ^53^ and only used to visualize the whole dataset with UMAP. *Cohort 1 and cohort 2 samples* were first normalized independently using the SCTransform function in Seurat V3 with parameters to regress out the following variables: experiment date, percent of mitochondrial gene mapping and the number of counts of each single cell (SCTransform(vars.to.regress = c(“experiment_date”,“percent_mito”, “nCount_RNA”), variable.features.n = 500). Integration of both cohorts together was then performed using the top 500 variable genes in both cohorts. Each cohort was projected into a common CCA (Canonical Correlation Analysis) space. Anchors (robust pairwise cell correspondence between datasets) were found by knn and snn graphs (FindIntegrationAnchors function) and they were used to transform the data in an integrated space. We used this integrated space, reduced to the top 40 PCs by PCA to visualize with UMAP ^72^. The number of significant principal components (PCs) was determined by comparison to PCA over a randomized matrix, as described by Klein et al. ^73^. Of note, another batch correction method, scAlign ^74^ produced similar results.

#### Clustering

We found shared clusters across samples using the Integration method in Seurat V3 as described in Stuart et al. ^53^. This time, *each sample* was first normalized independently using the SCTransform function in Seurat V3 with parameters to regress out the following variables: percent of mitochondrial gene mapping and the number of counts of each single cell (SCTransform(vars.to.regress = c(“percent_mito”, “nCount_RNA”), variable.features.n = 500). Integration of samples together was then performed using the top 500 variable genes. Each cohort was projected into a common reciprocal PCA space. Anchors (robust pairwise cell correspondence between samples) were found by knn and snn graphs (FindIntegrationAnchors) and they were used to transform the data in a shared space. We used this shared space and reduced it to the top 32 most significant PCs by PCA for clustering ^72^. In this space, a shared nearest neighbor graph was constructed from a k-nearest-neighbor graph (k = 20) by pruning cell-cell edges with less than 1/15 neighbor overlap. Community detection using the Louvain algorithm at a resolution of 0.5 found 11 clusters. Automated annotation using singleR with the BluePrintEncode reference data and manual annotation using canonical markers (see figure 3, 4, S7) clearly distinguished resting Tconvs, activated Tconvs, and Tregs. Type A IPEX Tregs were defined as Tregs in IPEX samples that overlapped with the HD Tregs (top left quadrant in UMAP). Type B IPEX Tregs are all the others.

#### Differential gene expression

We used limma-trend ^75^, as benchmarked in ^76^. Briefly, default TMM normalization from edgeR was applied to the SCTransform corrected count matrix. A linear model was fitted to the data using a contrast matrix including confounding variables (*lmFit* function). In order to find markers common across samples, or markers between type A and B IPEX Tregs, the contrast matrix contained the sample origin as a confounding variable. Empirical Bayes method was then used to estimate the overall trend of gene expression variance and adjust the genewise-wise residual variances towards this global trend (less variance for genes trending high and more for those trending low) (*eBayes(trend = TRUE, robust = TRUE)*). The *TopTable* function was then used to extract the differential statistics corresponding to the contrast of interest. Benjamin-Hochberg correction was applied to control type I error. Adjusted p-value < 0.05 were deemed significant.

### scRNAseq profiling of CD4+ T splenocytes cells in mice and bone marrow chimeras

scRNAseq profiling of mouse samples was performed in two experiments (see also Table S4). In the first experiment, 2 WT mice and 2 Δ*Foxp3* mice were profiled with the 10X Genomics Single Cell 3′ Reagent Kit (V2 chemistry), one channel per mouse. In the second experiment, CD4+ T cells from 3 bone marrow chimeras, 2 WT mice and 2 Δ*Foxp3* mice were pooled with sorted WT Tregs and Tconvs as controls and profiled with the 10X Genomics Single Cell 3′ Reagent Kit (V3 chemistry). Both experiments were analyzed together, after batch correction.

#### scRNAseq libraries

Spleens were harvested. After red blood cell lysis with ACK for 1 min at 4°C, 30% of splenocytes (∼30 million cells) were stained for sorting by flow cytometry in 200 uL of FACS buffer (PBS, 2% BSA) for 20 min at 4°C in the dark, with the following antibodies (1/100 dilution): 100 uL of FcBlock (homemade), TCRβ PE-Cy7 (H57-597; BioLegend), CD4 PerCPCy5.5 (GK1.5; BioLegend), DAPI. Whole Δ*Foxp3* and WT mouse CD4+ cells were sorted as DAPI^-^TCRb^-^CD4^+^. For BMC mice, Δ*Foxp3* cells were sorted as DAPI^-^TCRb^-^CD4^+^*CD45.1^+^CD45.2^+^*. WT cells were sorted as DAPI^-^TCRb^-^CD4^+^CD45.1*^+.^* 40-50,000 cells were single sorted in 50 uL of PBS-BSA 0.05% and the total volume was adjusted to a concentration of 1,000 cells/uL after cell counting with a hemocytometer. In the first experiment, cells from each tube were encapsulated in one channel of a 10x Chromium instrument, and libraries were constructed with a Single Cell 3′ Reagent Kit (V2 chemistry) (https://support.10xgenomics.com/single-cell-gene-expression/libraryprep/). In the second experiment, in order to pool samples for cell hashtagging, cells were sorted in 50 uL of PBS-BSA 0.05% in two different tubes (5,000 to 10,000 sorted cells for each sample) for sequencing in two 10X lanes, and libraries were constructed with a Single Cell 3′ Reagent Kit (V3 chemistry) (see also Table S4).

Libraries were sequenced on the NextSeq 500 platform (28/8/0/91, Read1/i7/i5/Read2). Gene counts were obtained by aligning reads to the mm10 transcriptome using CellRanger software (v3.0.2) (10X Genomics) (default parameters). The mm10 transcriptome was complemented with the transgenes sequences of the *Ires-Gfp*, *Yfp*, *Cre* in order to map reads to the *Foxp3* locus in *Foxp3^ires-gfp^*/B6 and *Foxp3^ΔGFPiCre^xR26^-YFP^* /B6 mice. In *Foxp3^ires-gfp^*/B6 mice, *Foxp3* locus expression was calculated as the sum of reads mapping to *Foxp3* and *Ires-Gfp*. In *Foxp3^ΔGFPiCre^xR26^-YFP^* /B6 mice, *Foxp3* locus expression was calculated as the sum of reads of mapping to *Foxp3* and *Cre* (not GFP because the Read 2 length is not long enough to reach the *GFP* sequence).

#### Hashtag libraries

See similar section in the human analysis (scRNAseq analysis of human PBMC samples)

#### Quality control

See similar section in the human analysis (scRNAseq analysis of human PBMC samples). SingleR was used with the Immgen reference data.

#### Batch correction

Experiment 1 and 2 were profiled with 10X V2 and V3, respectively. Batch correction was performed using the Integration method in Seurat V3 as described in Stuart et al. ^53^ and only used to visualize the whole mouse dataset with UMAP. Experiment 1 (10x V2) and experiment 2 (10X V3) samples were first normalized independently using the SCTransform function in Seurat V3 with parameters to regress out the following variables: percent of mitochondrial gene mapping and the number of counts of each single cell (SCTransform(vars.to.regress = c(“percent_mito”, “nCount_RNA”), variable.features.n = 500). Integration of both experiments together was then performed using the top 500 variable genes in both cohorts. Each cohort was projected into a common CCA space. Anchors were found by knn and snn graphs (FindIntegrationAnchors) and they were used to transform the data in an integrated space. We used this integrated space, reduced to the top 50 PCs by PCA to visualize with UMAP. The number of significant principal components (PCs) was determined by comparison to PCA over a randomized matrix, as described by Klein et al. ^73^.

#### Clustering

We found shared clusters across samples using the Integration method in Seurat V3 as described in Stuart et al. ^53^. This time, each sample was first normalized independently using the SCTransform function in Seurat V3 with parameters to regress out the following variables: percent of mitochondrial gene mapping and the number of counts of each single cell (SCTransform(vars.to.regress = c(“percent_mito”, “nCount_RNA”), variable.features.n = 500). Integration of samples together was then performed using the top 500 variable genes. Each cohort was projected into a common CCA space. Anchors were found by knn and snn graphs (FindIntegrationAnchors) and they were used to transform the data in a shared space. We used this shared space and reduced it to the top 28 most significant PCs by PCA for clustering. In this space, a shared nearest neighbor graph was constructed from a k-nearest-neighbor graph (k = 20) by pruning cell-cell edges with less than 1/15 neighbor overlap. Community detection using the Louvain algorithm at a resolution of 1.5 found 11 clusters. Automated annotation using singleR with the Immgen reference data and manual annotation using canonical markers (see Fig. S9) clearly distinguished resting Tconvs, activated Tconvs, resting Tregs and activated Tregs. Type A IPEX Tregs were defined as Tregs in IPEX samples that overlapped with the HD Tregs. Type B IPEX Tregs are all the others.

#### Differential gene expression

See similar section in the human analysis (scRNAseq analysis of human PBMC samples).

### Flow cytometry analysis of mouse cells

Viability dye (eFluor506, Thermo Fisher, dilution 1/1000) and antibodies against mouse CD4 BV-605 (clone RM4-5, Biolegend, dilution 1/500), Thy1.2 APC-Cy7 (clone 30-H12, Biolegend, dilution 1/500), CD25 PE-Cy7 (clone PC61, Biolegend, dilution 1/500), CD45.1 PE-Cy7 (clone A20, Biolegend, 1/500), CD45.2 A700 (clone 104, Biolegend, 1/500), were used. Cell suspensions were stained for surface markers and viability dye for 20 min on PBS/0.5%FCS. Data acquisition was performed on a BD Fortessa cytometer using DIVA software (BD Biosystems) and analyzed using FlowJo (Tree Star).

### Statistics

Analysis was done using R-3.6.2. Heatmaps were made with Morpheus (https://software.broadinstitute.org/morpheus) or the pheatmap package in R (https://cran.r-project.org/web/packages/pheatmap/index.html). All other plots are made with ggplot2 ^77^. Statistical tests are described in their respective method section.

## SUPPLEMENTARY FIGURE LEGENDS

**Fig. S1.**
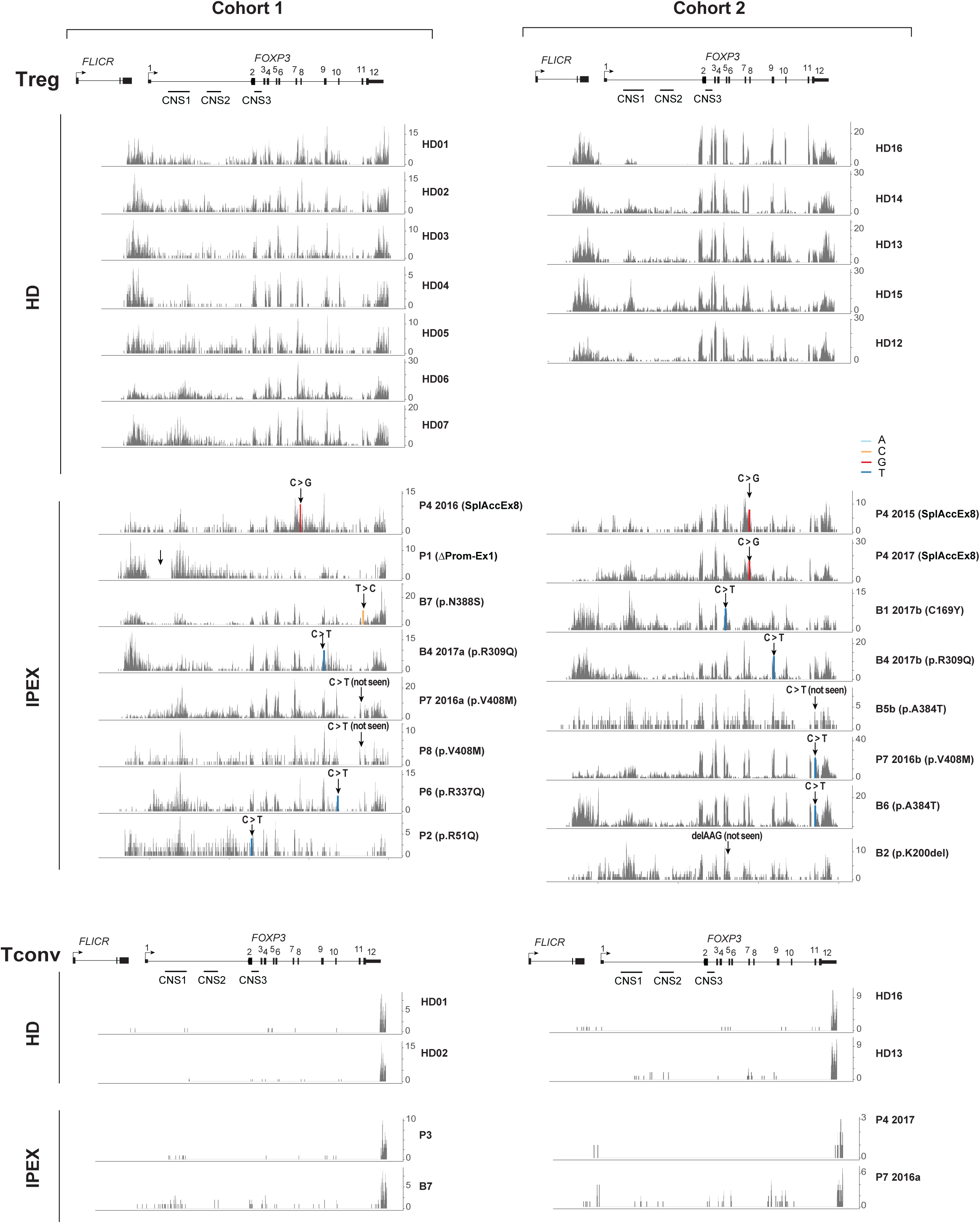
*FOXP3* locus activity in IPEX Treg-like cells. Read mapping to the FOXP3 locus from sorted CD4+ CD25+ CD127low cells in HD and IPEX samples from both cohorts (population RNAseq). Arrow indicates mutation. Mutated bases are colored. All samples from both cohorts are shown as well as control traces from representative Tconvs samples (sorted CD4+ CD25- CD127+ cells)

**Fig. S2.**
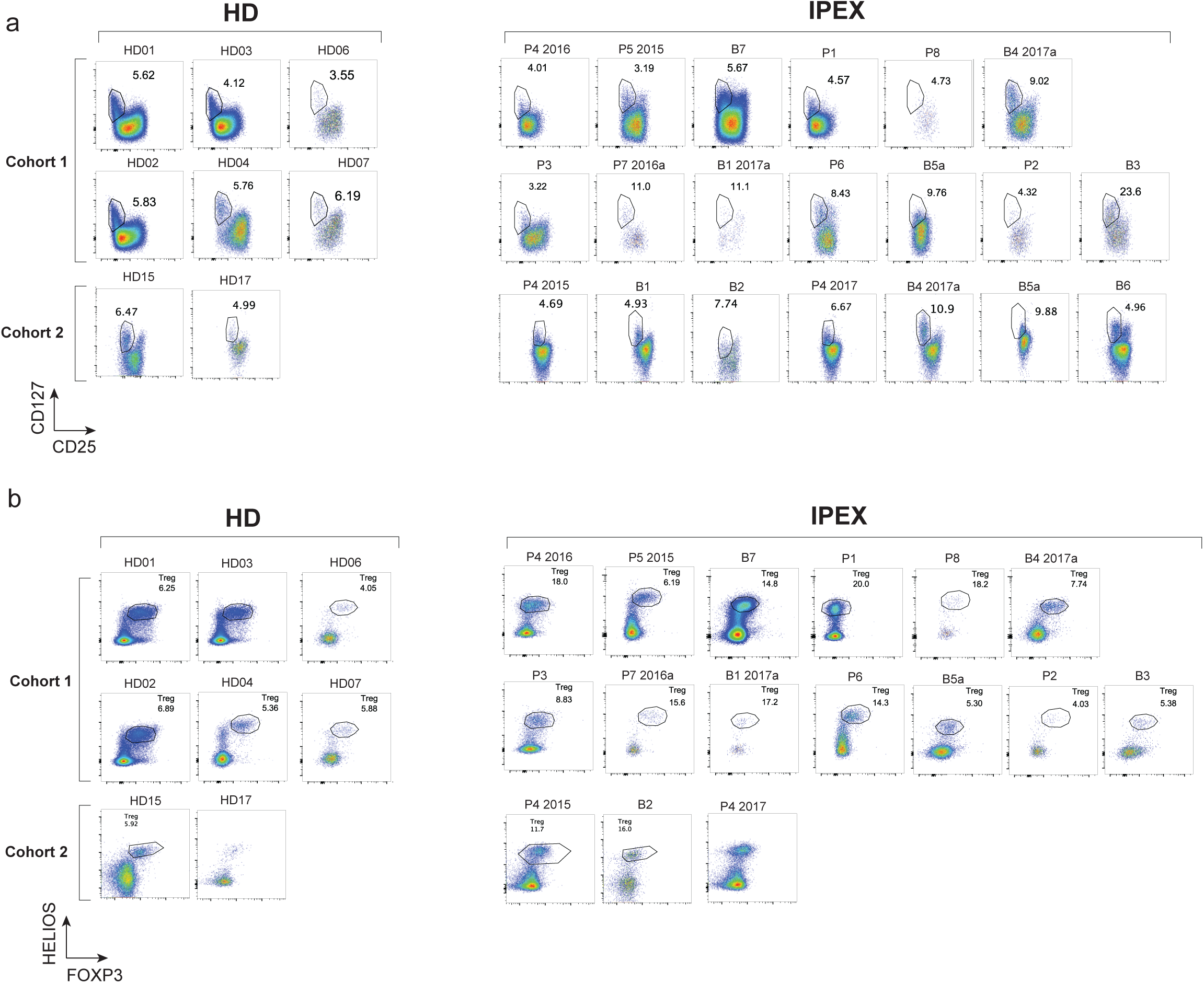
Flow cytometric analysis of Treg-like cells in IPEX. **a, b.** Flow cytometric analysis of HD and IPEX CD4+ cells: CD25 and CD127 (a), FOXP3 and HELIOS (b). Gated cells are HD Tregs and IPEX Treg-like cells. All samples from both cohorts are shown.

**Fig. S3.**
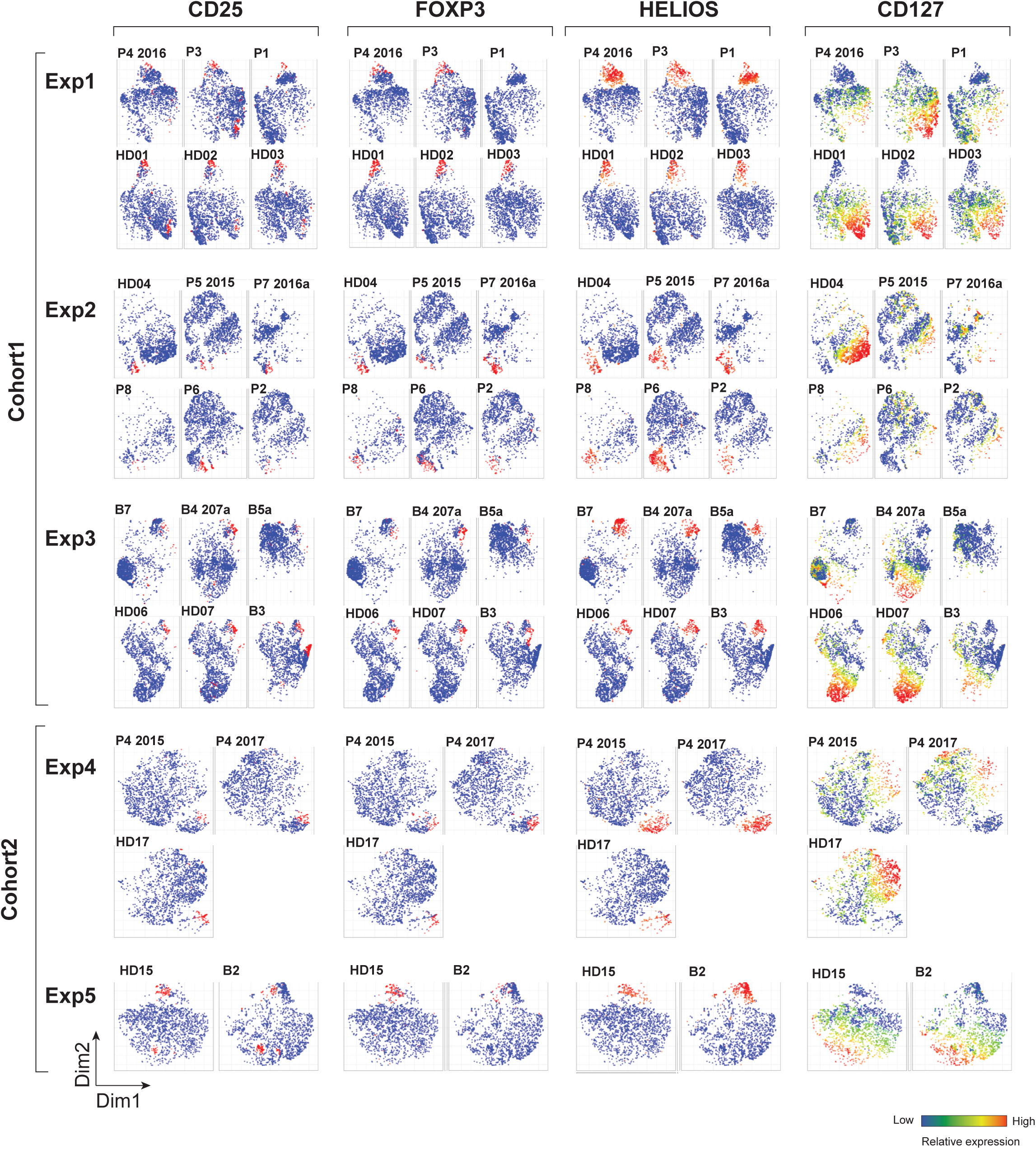
Identification of Treg-like cells in IPEX by flow-tSNE. Flow-tSNE plots of CD3+ CD4+ cells using flow cytometric expression of CD3, CD4, CD25, CD127, HELIOS, CD45RA and FOXP3. Color represents scaled expression of CD25, FOXP3, HELIOS and CD127. All samples from both cohorts are shown.

**Fig. S4.**
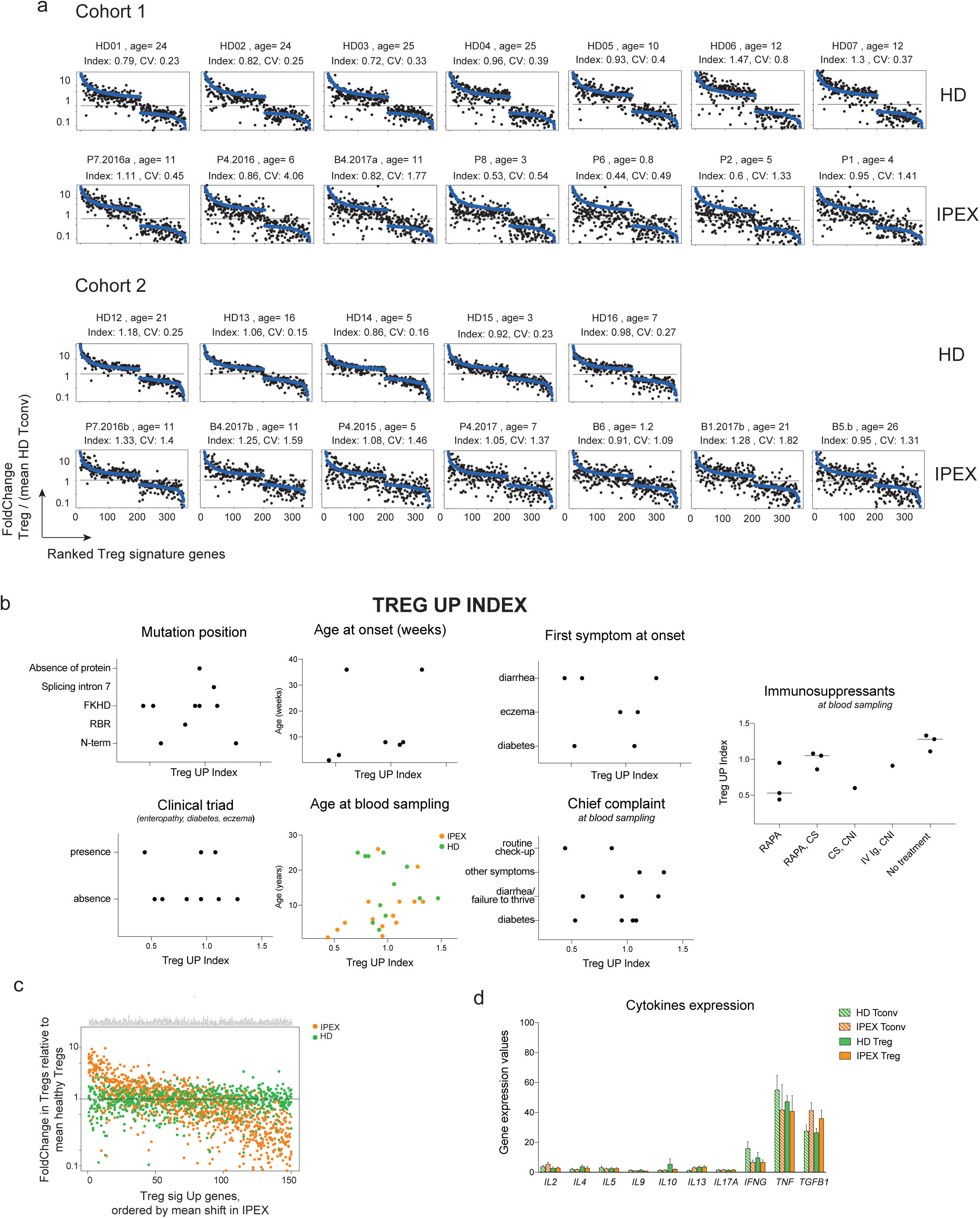
IPEX Treg-like cells maintain expression of the Treg signature, but with increased noise. **a.** Same ranked FC plots as 2b. showing the distribution of each Treg signature gene expression in IPEX and HD donors. The y-axis displays the gene expression foldchange in each donor over the average expression in HD Tconvs. Genes are ranked by the average Treg over Tconv foldchange in HD. Regression lines are shown in blue. Age (in years) and summary statistics (intensity (index) and variability (coefficient of variation, CV) of expression the Treg signature), are shown for each sample. **b.** Clinical correlations with the Treg UP index (computed in a). RAPA, rapamycin, CS, corticosteroids, CNI, calcineurin inhibitor, IV Ig, intravenous immunoglobulins. **c.** Ranked FC plot showing the distribution of expression of the Treg signature gene in IPEX (orange) and HD (green) Treg samples. The y-axis corresponds to the expression ratio in IPEX Treg-like cells relative to the mean in HD Tregs. Genes are ranked by their average shift in IPEX samples. **d.** Average gene expression of cytokines across Treg and Tconvs in IPEX and in HD (bar plot with mean and SEM, normalized counts).

**Fig. S5.**
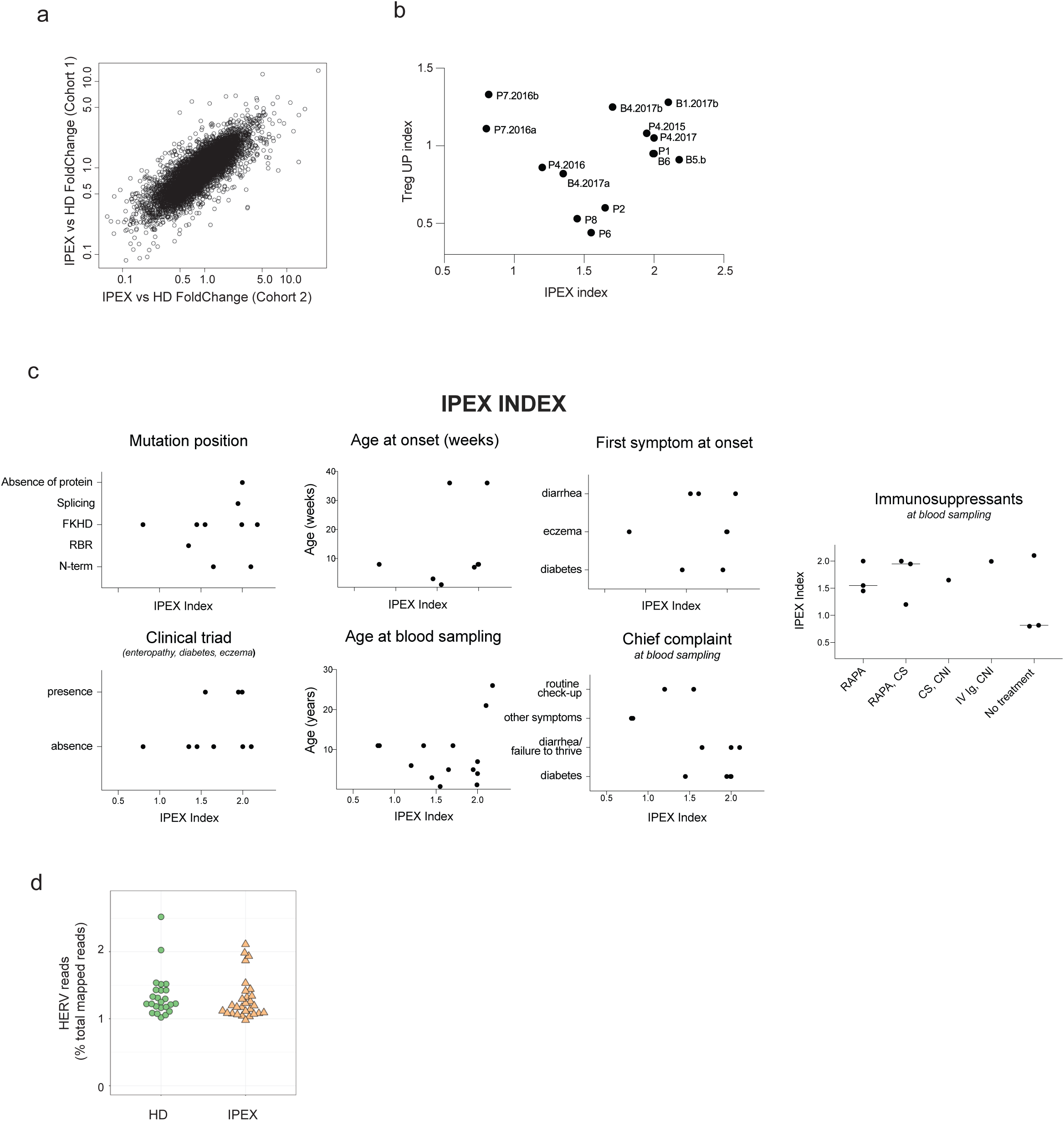
IPEX signature: reproducibility, independence from the Treg up index, and clinical correlates. **a.** IPEX/HD expression ratio in cohort 1 vs cohort 2 showing the reproducibility of the IPEX effect between the two cohorts. **b.** Absence of correlation between the IPEX index (x-axis) and Treg UP index (y-axis) **c.** Clinical correlations with the IPEX index. RAPA, rapamycin, CS, corticosteroids, CNI, calcineurin inhibitor, IV Ig, intravenous immunoglobulins. **d.** Proportion of human endogenous retroviruses (HERV) mapped reads in IPEX and HD.

**Fig. S6.**
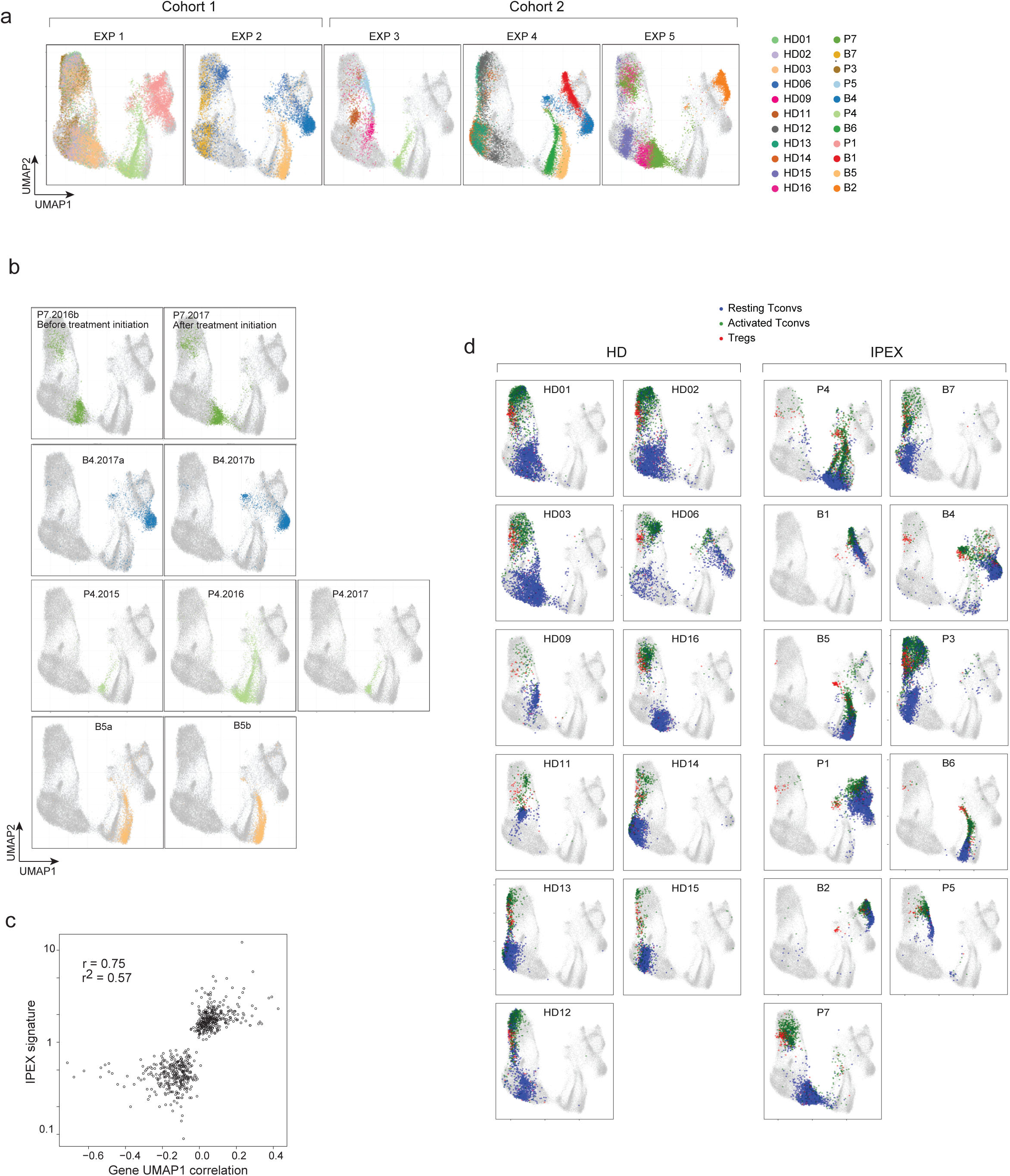
scRNAseq analysis of CD4+ cells in IPEX and HD reveals a stable IPEX signature that affect all CD4+ cells (resting, activated Tconvs and Tregs) **a.** Same UMAP plots as 3a., showing the reproducible segregation of HD and IPEX samples: plots are split by experiment and cohort. Individual HD and IPEX donors are highlighted in different colors. **b.** Same UMAP as 3a. showing the stability of the IPEX transcriptomic signature: before and after treatment initiation (P7), over several days (B4a and B4b, four days apart) and or years (> 2 years apart, P4), and across two different experiments (B5, technical replicates) **c.** UMAP1 correlates with the IPEX signature defined by population RNAseq (Pearson correlation r = 0.75). y-axis shows the IPEX/HD expression ratio of the IPEX signature genes (population RNAseq). The x axis shows their correlation with UMAP1 in scRNAseq. **d.** Same UMAP as 3a. showing each HD and IPEX donor individually. Blue, green, and red cells represent resting Tconvs, activated Tconvs, and Tregs, respectively.

**Fig. S7.**
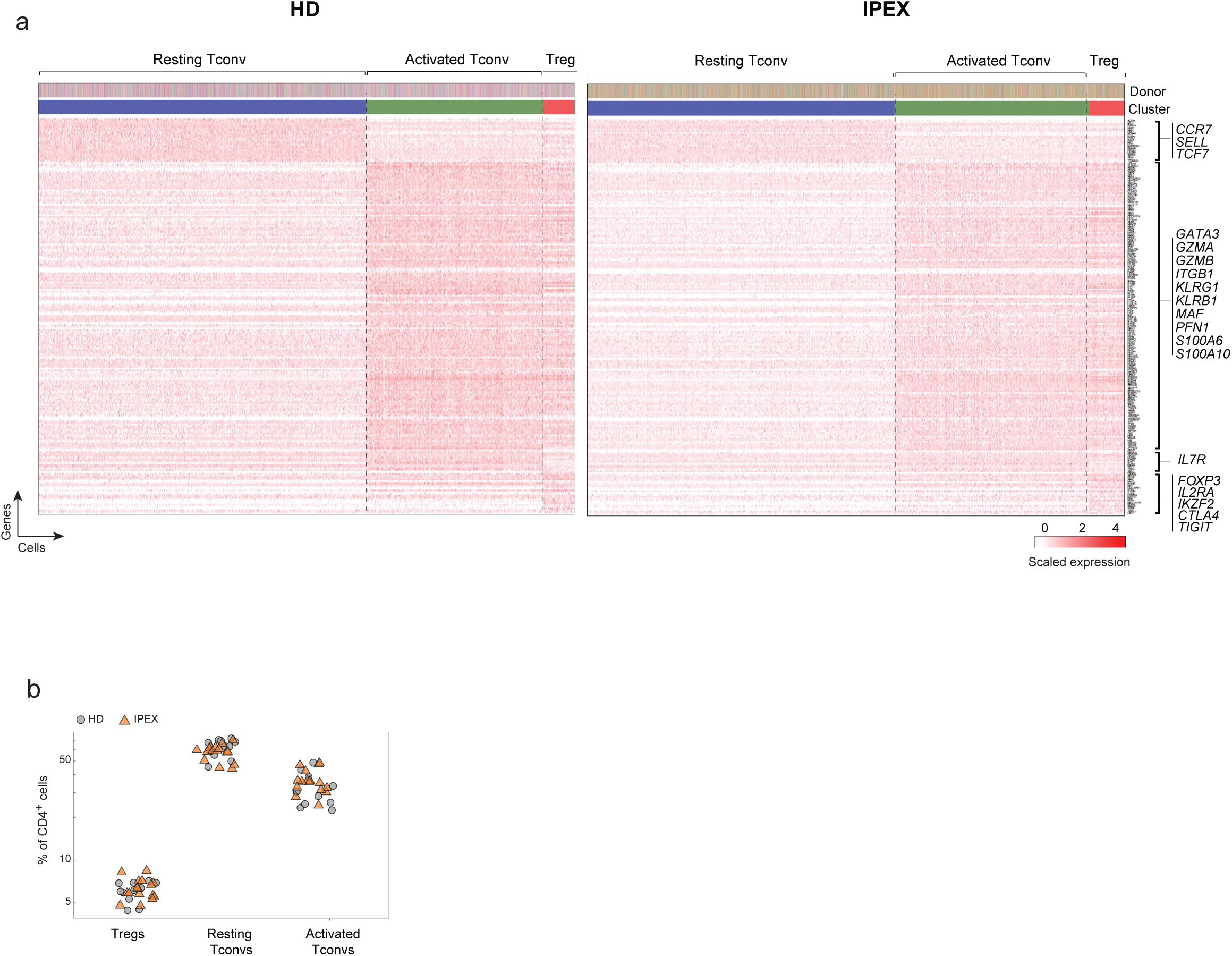
Identifcation of resting Tconvs, activated Tconvs and Tregs in IPEX by scRNAseq. **a.** Single-cell biclustering heatmap of canonical resting Tconv, activated Tconv and Treg genes. Top ribbons indicate donor origin and annotations for every single cell. **b.** Similar proportions of Tregs, resting and activated Tconvs in total CD4+ cells in HD and IPEX (scRNAseq).

**Fig. S8.**
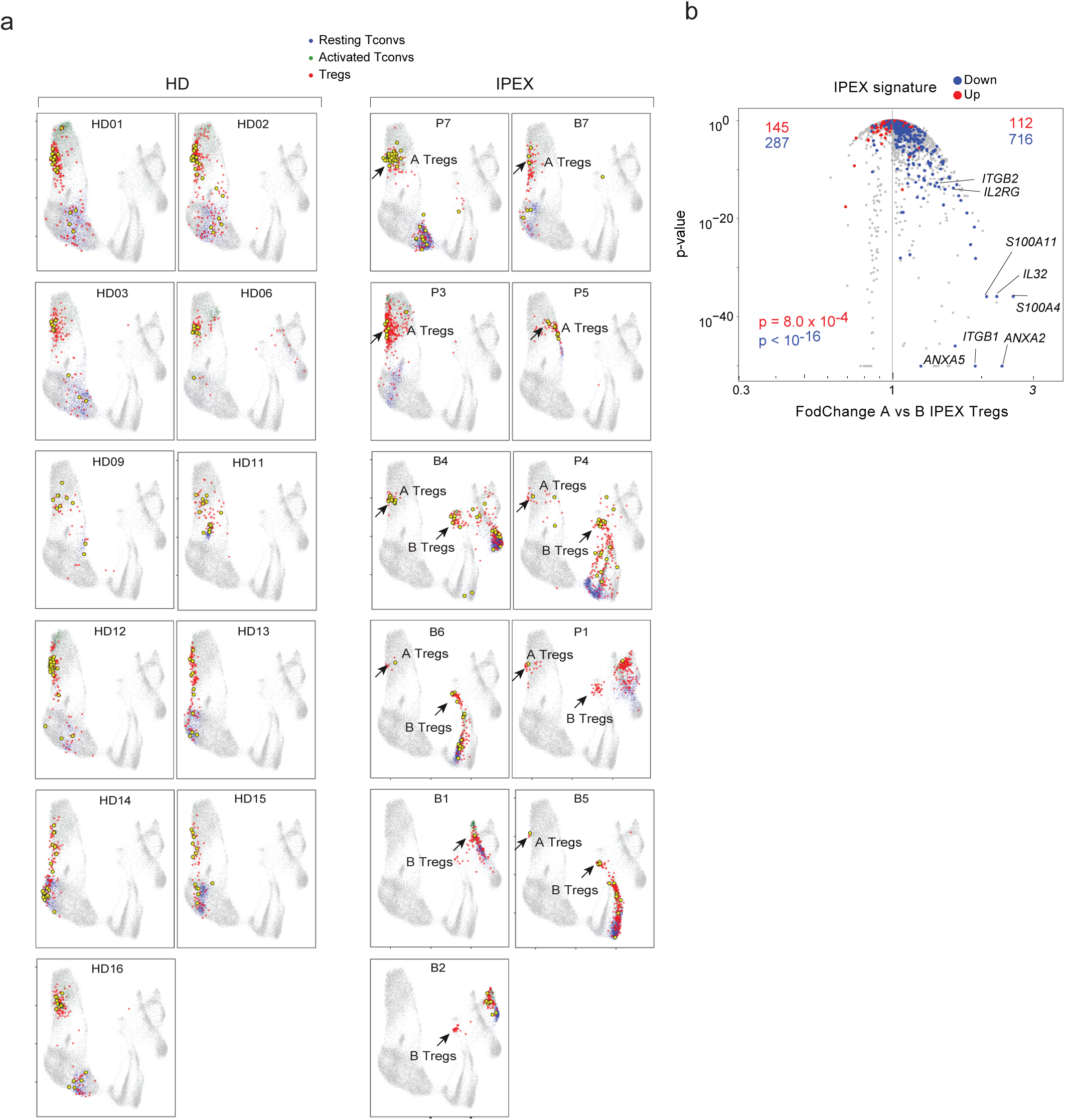
Heterogeneous Treg-like cells in IPEX (A and B types) identified by scRNAseq. **a.** Same UMAP as 3a. showing each HD and IPEX donor individually. Blue, green, and red cells represent resting Tconvs, activated Tconvs, and Tregs, respectively. *FOXP3-*expressing cells (RNA) are in yellow. An arrow indicates type-A IPEX Tregs, overlapping with HD Tregs. **b.** Down-tuning of the IPEX signature in type A IPEX Tregs vs. type B IPEX Tregs. Volcano plot comparing the gene expression profiles of type A versus type B IPEX Tregs. Up- and downregulated signature genes are highlighted (red and blue, respectively). χ2 -test p values.

**Fig. S9.**
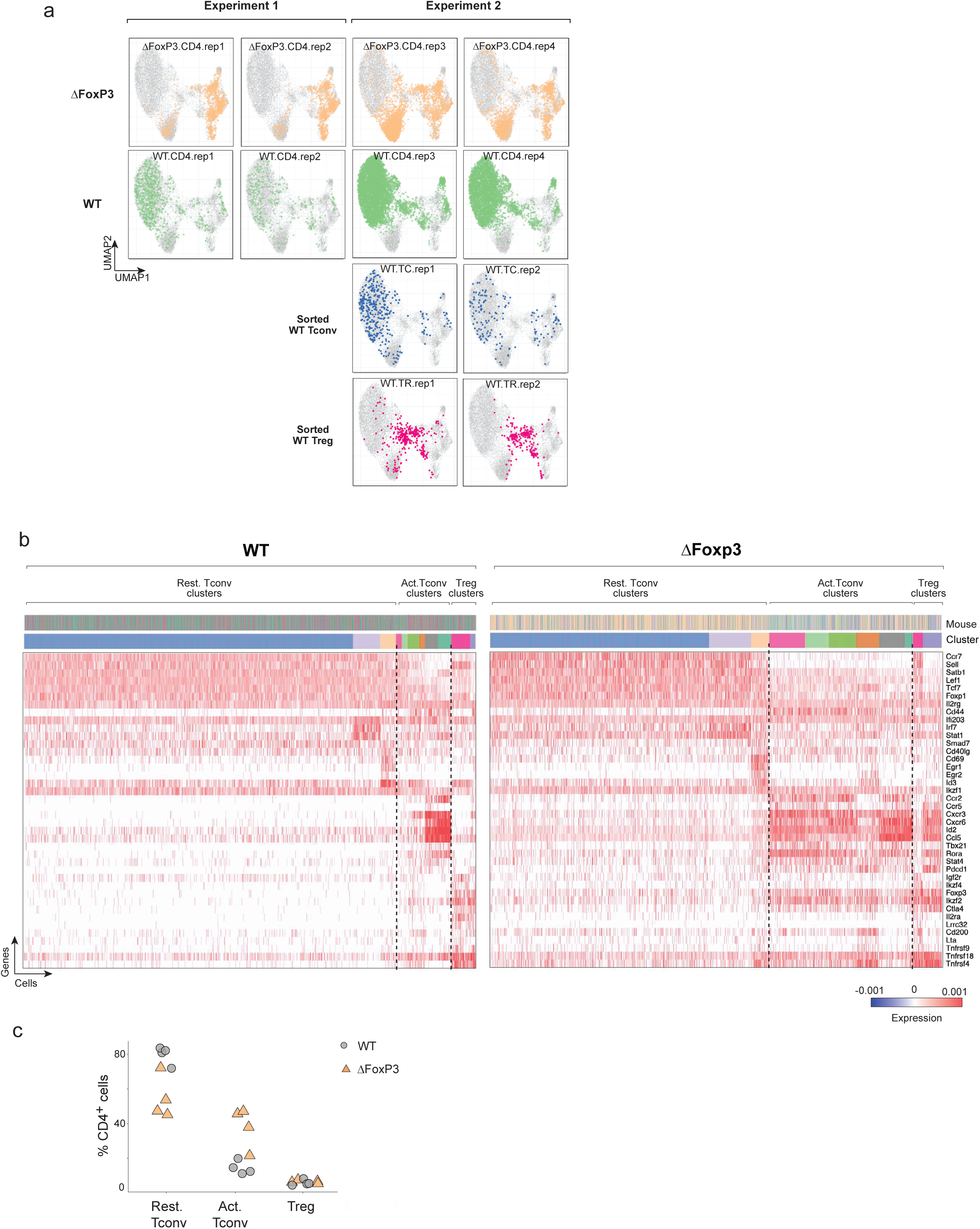
Identifcation of resting Tconvs, activated Tconvs and Tregs in Δ*Foxp3* mice by scRNAseq. **a.** Same UMAP as 5a showing the individual samples in both experiments and the absence of batch effect. **b.** Single-cell biclustering heatmap of the expression of canonical genes in WT and Δ*Foxp3* Tregs, resting Tconvs, and activated Tconvs. Top ribbons indicate mouse origin and cluster annotations for each single cell. **c.** Proportion of Tregs, resting and activated Tconvs in total CD4+ cells in WT and Δ*Foxp3* mice (scRNAseq).

**Fig. S10.**
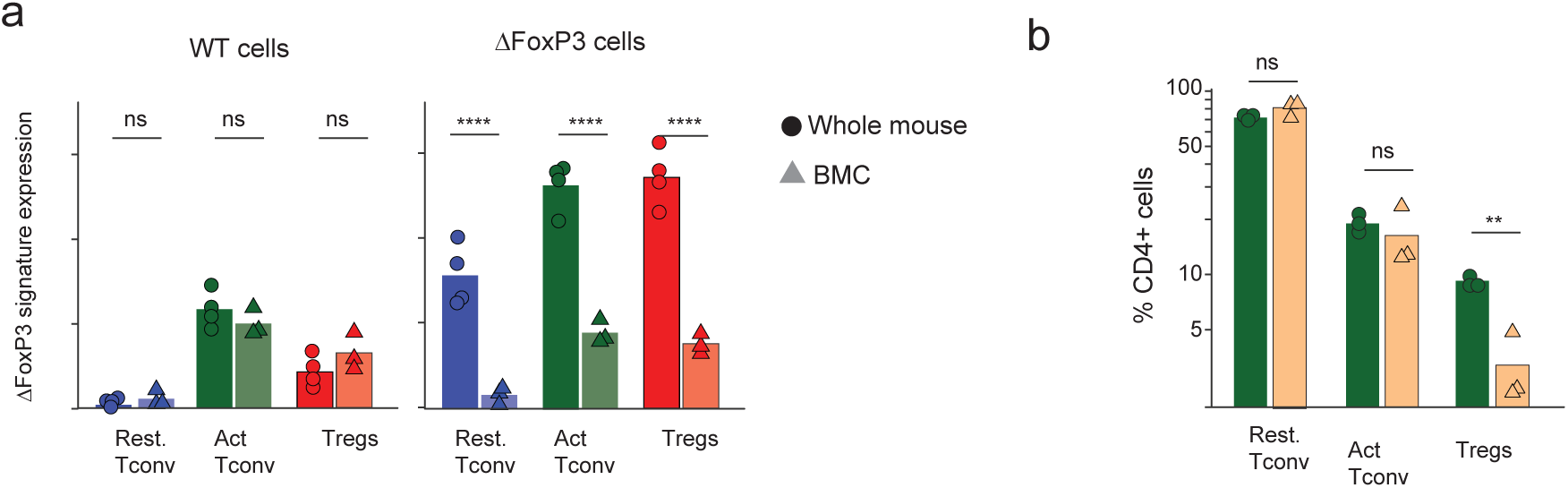
Lower proportion of Δ*Foxp3* Tregs and absence of the Δ*Foxp3* signature in **Δ***Foxp3* Tconvs and Tregs in mixed bone marrow chimera with WT cells. **a.** Δ*Foxp3* signature expression in whole mice and mixed bone marrow chimera (50/50 WT and Δ*Foxp3*) showing the downregulation of the Δ*Foxp3* signature expression in resting, activated Δ*Foxp3* Tconv and Δ*Foxp3* Tregs in 50/50 BMC. **b.** WT Tregs outcompete Δ*Foxp3* Tregs in in 50/50 BMC. Proportion of WT and Δ*Foxp3* Tregs, resting and activated Tconvs in total CD4+ cells.

## SUPPLEMENTARY TABLE LEGENDS

**Supplementary Table 1. Clinical characteristics of healthy donors (HD) and IPEX patients by cohort (summary table and singular values)**

**Supplementary Table 2. IPEX signature genes with their average IPEX/HD fold change in Tregs and Tconvs.**

**Supplementary Table 3. CD4+ signatures significantly enriched in the IPEX signature** (p<0.001, hypergeometric test, extracted from the ImmuneSigDB or manually extracted by curation of CD4+ relevant published dataset in human)

**Supplementary Table 4. List of human and mouse scRNAseq datasets (human and mice) with quality controls metrics**

